# Correcting and georeferencing natural history occurrence records with calibrated uncertainty

**DOI:** 10.64898/2026.10.01.755955

**Authors:** Ann M Raiho, Emmy Spadaro, Amanda Tieu, Wilson Guillory, James Ackerman, Zhe Zhu, Zhiqiang Yang, S Blair Hedges

## Abstract

Occurrence records underpin biodiversity analyses, but their coordinates are often erroneous or missing. Misplaced coordinates distort environmental predictors and range estimates used in conservation assessment. Existing tools flag suspect records or georeference locality text, but rarely correct coordinates, provide calibrated uncertainty, or quantify downstream consequences.
We present GeoMend, a deterministic, rule-based method that flags and corrects suspect coordinates and georeferences records lacking coordinates from locality text, assigning each a precision tier or uncertainty radius. It uses no machine learning or artificial intelligence. We evaluate it against expert coordinates for 16,913 records and 736 species on Hispaniola, compare georeferencers using 5,000 localities from 72 countries, and measure changes in environmental predictors and the extent of occurrence.
On held-out Hispaniola records, GeoMend flags 92% of coordinates more than 5 km from expert truth, versus 27% for CoordinateCleaner. Among the accepted corrections with paired coordinates, the median error falls from 2.19 to 0.82 km, and 68% of the complete held-out pairs improve. On records placed by both GeoMend and GEOLocate that also have original coordinates, the median error is 0.67 km for GeoMend versus 10.69 km for GEOLocate. For records lacking coordinates, evaluated uncertainty radii contain expert truth about 92% of the time in well-supported gazetteer classes. Correction reduces the mean standardized environmental distance from expert truth by 38% and moves the extent of occurrence toward expert truth by a median of 8% (mean of 79%). Corrections and newly georeferenced records give 17 Hispaniola species their first range estimate.
On the global benchmark, GeoMend is more accurate than GPT-4o for two-thirds of the records placed by all four comparators, with median errors of 4.4 and 10.8 km, respectively. Applied to Global Biodiversity Information Facility (GBIF) records lacking coordinates, it places records from 82,180 previously unmapped species across 233 countries, ranging from precise locality interpretations to administrative centroids. Automated placement makes the global backlog more approachable, while explicit tiers distinguish uses supported by precise and coarse coordinates. Further georeferencing requires regional expertise to combine automated placement with local knowledge.

**Data and code for peer review:** The GeoMend source code, its configuration, and the derived data tables underlying every figure and number in this paper are available to reviewers as an anonymised archive, through an anonymous view-only link that hides contributor names (https://osf.io/mcdhb/?view_only=ab3de27644ed437994d1cf3c30da079b). On acceptance, the code and data will be deposited in a permanent public repository and the DOI will be added to the Data Availability Statement on the title page.

## 1. Introduction

Preventing species extinctions depends on identifying where conservation action is needed, which requires reliable knowledge of species distributions and how they may change (Guisan et al., 2013; Margules & Pressey, 2000). A species distribution model reads environmental predictors at the occurrence coordinates (Elith et al., 2011), and the size and location of a species’ range inform International Union for Conservation of Nature (IUCN) Red List assessments (IUCN, 2012). Both uses rely on accurate coordinates representing habitats where specimens were collected.

Natural history collections hold the deepest record of these occurrences. A survey of 73 large natural history museums in 28 countries documented 1.1 billion specimens (Ariño, 2010; Johnson et al., 2023) where many predate Global Positioning System (GPS) coordinates (Stenhouse et al., 2025). These records are aggregated by the Global Biodiversity Information Facility (GBIF) (Shirey et al., 2019). In the complete 1 July 2026 GBIF preserved-specimen snapshot, more than 410,000 species, or about 28% of all species represented, have no coordinate on any record (GBIF.org, 2026a). This large fraction of Earth’s species is therefore absent from analyses that require georeferenced occurrences, with the gap falling disproportionately on older specimens and under-collected regions (Hortal et al., 2015; Meyer et al., 2015).

Converting this backlog into mapped occurrences manually is slow and error-prone (Barber et al., 2013; Blagoderov et al., 2012; Vollmar et al., 2010). In manual georeferencing, a human researcher translates a written locality such as “10 km SW of San Juan” into a coordinate. This is typically done using a map or gazetteer to locate the locality, then locating the position of the occurrence using contextual clues such as roads or landmarks, but doing so manually takes minutes to hours per record. Errors arise when a stated distance is drawn in a straight line instead of following along a road, when a transcription is wrong, or when language, place names, and administrative boundaries have changed since collection (Chapman & Wieczorek, 2020; Wieczorek et al., 2004).

Occurrence records contain both positional errors, described in the previous paragraph, and sampling biases, which arise because collectors visit some places, such as roadsides and accessible habitats, more often than others (Daru et al., 2018; Kadmon et al., 2004). Incorrect coordinates for either reason misrepresent the environmental conditions where specimens were collected and can inflate or shrink estimated ranges (Gábor et al., 2022; Graham et al., 2008; Mitchell et al., 2017). The tool presented here, GeoMend, addresses positional errors and not sampling bias by correcting where records are mapped using locality text and context information.

Within positional errors, there are many potential causes, including coordinates placed along the wrong part of a road, mismatches between coordinates and locality text, false precision, and locations that conflict with a species’ known range or environment. Existing tools address only part of the issue. Coordinate cleaners flag but do not georeference. For example, CoordinateCleaner flags coordinates that fall in the ocean, at country or province centroids, or at known biodiversity institutions (Zizka et al., 2019), bdc adds checks on non-spatial fields such as names and dates (Ribeiro et al., 2022), and broader packages wrap these steps into acquire-and-clean workflows (Jin & Yang, 2020; Patten et al., 2024). A separate family of tools produces a coordinate from a written locality. GEOLocate and recent large-language-model approaches parse locality text (Rios & Bart, n.d.; Xie et al., 2025), and HOGS reads historical maps (Ravenscroft et al., 2025). These tools georeference records that lack a coordinate but do not correct an existing incorrect coordinate, and most tools return a point with no calibrated statement of how uncertain it is. There are no available tools to flag, correct, georeference, and assign uncertainty in a single pass.

Here, we present GeoMend, an open, deterministic method that flags, corrects, and georeferences in one procedure, assigning each output a precision tier from road-level placement (∼1 km uncertainty) to administrative fallback (∼40 km uncertainty). GeoMend does not use artificial intelligence: its decisions follow explicit rules and reference data, so the same inputs and settings produce the same outputs.

Large language models can georeference locality text with accuracy comparable to established services (Fernando et al., 2026; Xie et al., 2025). However, general models typically provide neither calibrated uncertainty nor abstention when evidence is insufficient. Both are established georeferencing requirements (Chapman & Wieczorek, 2020; Wieczorek et al., 2004; Xie et al., 2025). We therefore compare GeoMend with a language model to assess accuracy and generalization, while treating reproducibility, uncertainty, and abstention as separate methodological design criteria. We evaluate detection and placement against expert truth on Hispaniola (Section 3.1), test global generalization against existing tools (Section 3.2), and measure downstream range recovery and the occurrence data returned for under-sampled species (Section 3.3).

## 2. Materials and Methods

### 2.1 The GeoMend method

GeoMend uses occurrence metadata to make one of four decisions: retain a coordinate, correct it, georeference a record that lacks one, or flag the record for review (Fig. 1a). Corrected coordinates enter the high-precision Tier 1a. Coordinate-less records may receive a structured-text placement (Tier 1b), a gazetteer centroid (Tier 2), or an administrative centroid (Tier 3). Existing coordinates that cannot be corrected reliably are either kept with elevated uncertainty or flagged.

**Figure 1.**
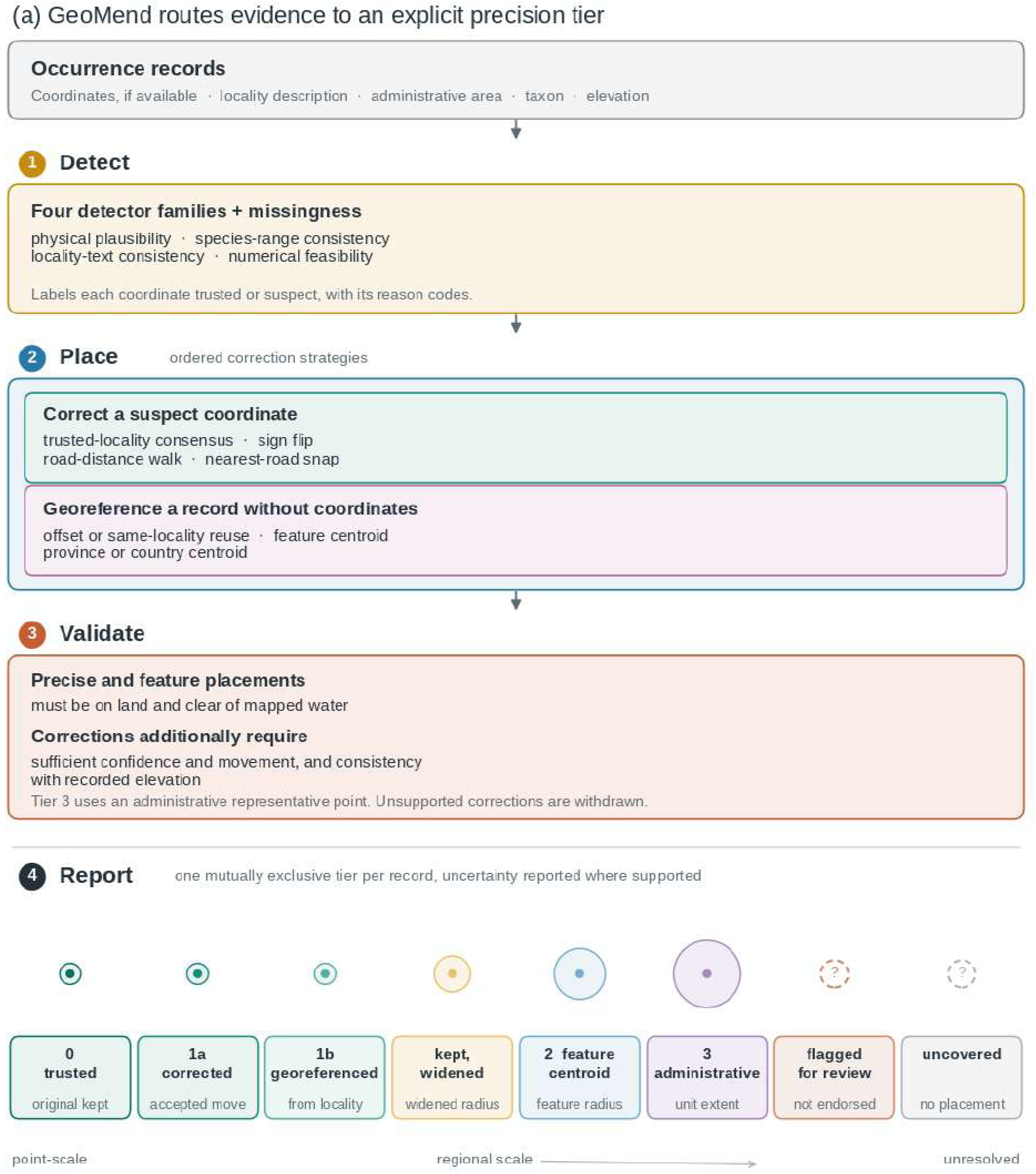
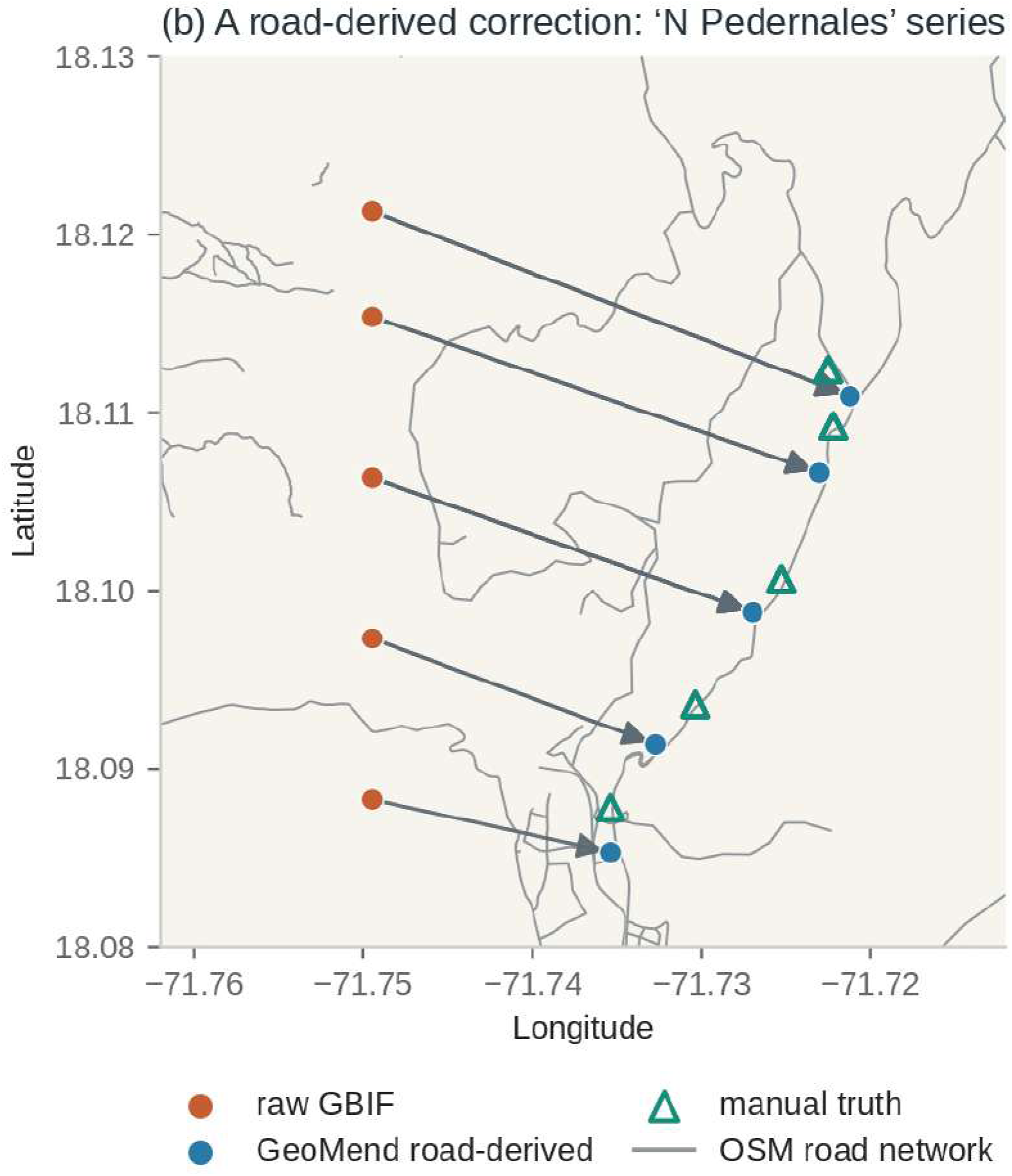
GeoMend’s workflow schematic and one worked correction. **(a)** Four operations summarize the implementation: four detector families plus coordinate missingness label the input, ordered strategies correct suspect coordinates or georeference records without coordinates, validation gates withdraw unsupported positions, and every record is reported in one mutually exclusive precision tier. **(b)** In the “N Pedernales” series, raw points follow an artificial straight-north line. Measuring the stated distance along the road and applying the 50 m habitat offset reduces median error from 2.63 to 0.27 km against held-out manual truth. The branch-level decision tree is retained as Fig. S2.

The method combines five regional reference layers: roads, water bodies, elevation, administrative boundaries, and a gazetteer. For Hispaniola, we used OpenStreetMap roads and water for Haiti and the Dominican Republic, a 30 m elevation model, GADM boundaries, and GeoNames (Geofabrik GmbH, n.d.; GeoNames, n.d.; OpenStreetMap contributors, n.d.). Applying GeoMend else-where requires the corresponding layers for the new region, but no part of the method depends on the Hispaniola files themselves. Its thresholds are explicit rules or summaries of manual coordinates, not fitted model parameters.

GeoMend’s central operation is to translate locality text into a position (Aldana-Bobadilla et al., 2020; Halterman, 2017). It parses a description such as “10 km SW of San Juan” into a distance, bearing, and anchor place, then follows the stated distance along the road network from that anchor rather than drawing a straight line. This road-walk can correct an inaccurate coordinate or place a record that lacks one (Fig. 1b).

Existing coordinates first pass through four detector families. These test physical plausibility, consistency with the species’ range, agreement with the parsed locality text, and numerical feasibility (Supplementary Methods S1). A record that fails any test passes through four correction strategies in order: consensus snapping, sign-flip repair, the road-distance walk, and nearest-road snapping. Off-trail taxa, defined a priori based on the field sampling design, are exempt from nearest-road snapping, and their road-distance corrections are capped at 5 km (S2). This movement cap limits large, potentially harmful road-based displacements.

Coordinate-less records are placed only as precisely as their locality text allows. A distance and bearing from a named place supports a road-walk (Tier 1b). A place name without an offset supports its gazetteer point (Tier 2). A province or country alone supports the centre of that unit, with the unit’s size as the uncertainty radius (Tier 3). A record without a locatable place remains unplaced.

Validation gates prevent unsupported placements from entering the output. Our implementation is currently focused on terrestrial species, so Tier 1b and Tier 2 points must fall on land and outside mapped inland water. A correction must also pass thresholds for confidence, minimum movement, and agreement with the elevation model. If a correction fails, GeoMend keeps the original coordinate with a wider uncertainty radius or sends the record to expert review when the text cannot support a repair (S3, S4).

Every GeoMend output states the precision that its evidence supports. Each tier has an error distribution measured against the expert coordinates, so users who retain only Tiers 0 to 1b can see the median error of the records they keep (Table S3). Coarse placements and unchanged coordinates retained with uncertainty also receive a record-level radius. For retained coordinates, the radius contains 90% of calibration errors for the same flag reason. For gazetteer placements, it contains 90% of errors in the same GeoNames feature class. For administrative placements, it is the size of the named unit. The precise tiers are summarized by their error distributions instead (Section 3.1.3, Supplementary Methods S2 and S4). The radius for GeoNames spot features remains provisional because no calibration record belonged to that class (S2).

### 2.2 Datasets

Three datasets serve distinct roles in development, validation, and global evaluation. The first provides expert ground truth: 16,913 manually curated and georeferenced records from Hispaniola, spanning 736 species of frogs, reptiles, and orchids. Over two years, the curators located specimens using Google Maps and Google Earth, contemporary and historical topographic maps, including the 1:50,000 CaribMap sheets for Haiti (1962) and the Dominican Republic (1988) (Hedges, 2013), online gazetteers including the NGA GEOnet Names Server, and collectors’ original records when available (National Geospatial-Intelligence Agency & U.S. Board on Geographic Names, n.d.). The expert retained record-level notes documenting each decision. The curated frog dataset combines GBIF-linked records (GBIF.org, 2026b) with unpublished personal-collection records supplied by the amphibian curator (Center for Biodiversity, Temple University, 2025). Species are grouped by the genus and species labels supplied in the curated tables. GeoMend does not independently reconcile taxonomic splits, instead revised assignments affect species-level reference ranges and EOO, while positional error is evaluated record by record.

The reptile records provide the best-documented example of how the Hispaniola dataset was curated, and we publish them in this paper. Their curation extended beyond assigning coordinates. The curator compared GBIF locality text with original paper records when available, resolved vague or missing identifications using the taxonomic literature, updated the names to current taxonomy, validated the records, and assembled the final dataset. Table S4 lists the taxonomic sources used. Separate curators georeferenced each taxonomic group: one expert curated amphibians, two curators curated reptiles, and another expert curated orchids. These curated coordinates and notes guided GeoMend’s development.

Placement accuracy is evaluated on 6,322 records (37%) withheld from threshold selection. This validation set comprises a random 20% of the frog and reptile records and every orchid record. Holding out orchids as an entire taxon, with coordinates assigned by a different expert, provides a stronger test of cross-taxon generalization.

We test the one remaining dependence on expert data rather than assuming that it is harmless. Range-consistency detectors build each species’ range from expert coordinates, including held-out records from that species. We therefore repeat the detection evaluation after rebuilding every reference range from GeoMend’s own first-pass output, so no expert coordinate enters the alternative reference (Section 3.1.1).

The second dataset tests geographic generalization. It is the published Nature Plants benchmark of 5,000 localities from 72 countries, which we use for direct comparison with other georeferencing methods (Section 2.6) (Xie et al., 2025).

The third dataset measures the global backlog that GeoMend could address. It consists of species with preserved specimens in GBIF but no coordinate on any record. We identified this backlog from the complete 2026-07-01 GBIF Parquet snapshot rather than from a pre-selected download (GBIF.org, 2026a). Of 1,491,014 species represented by preserved specimens, 410,649 (27.5%) had no coordinate on any record. Among these species, 262,794 had usable locality text. We georeferenced all 898,176 of their records from 245 countries.

Because backlog records lack coordinates, they cannot provide ground truth for their new placements. We therefore constructed a stand-in test set of 500,698 coordinate-bearing records, with reweighted sampling to mimic the distribution by coordinate-less records in clade, presence of a locality offset, and locality-description length. We ran GeoMend on their locality text and compared the resulting position with the coordinate already present. This comparison measures agreement with an existing georeference and gives a proxy for expected backlog accuracy (Supplementary Methods S5).

### 2.3 Calibration

GeoMend makes its decisions using 38 documented numeric thresholds. None is fitted as a model parameter. Instead, each threshold is measured from the expert coordinates, taken from a physical or field-practice quantity, or chosen by the authors and shown through sensitivity analysis not to affect the results (Table 1). Table S1 gives every value and its basis.

**Table 1.**
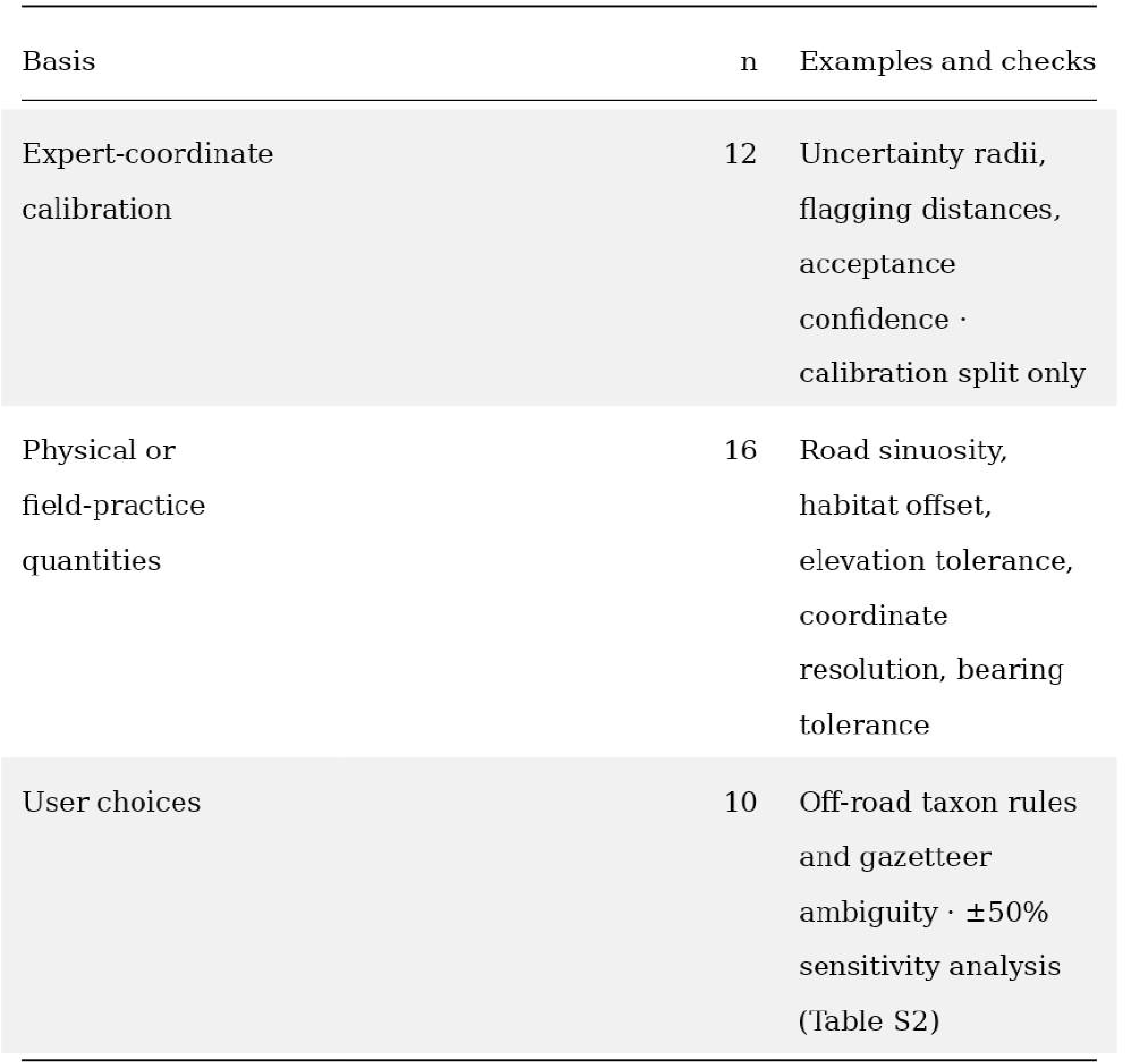
How each of GeoMend’s 38 flagging and coordinate-placement thresholds is set.

| Basis | n | Examples and checks |
| --- | --- | --- |
| Expert-coordinate calibration | 12 | Uncertainty radii, flagging distances, acceptance confidence · calibration split only |
| Physical or field-practice quantities | 16 | Road sinuosity, habitat offset, elevation tolerance, coordinate resolution, bearing tolerance |
| User choices | 10 | Off-road taxon rules and gazetteer ambiguity · $\pm 50\%$ sensitivity analysis (Table S2) |

Twelve thresholds are calibrated on expert coordinates without using the validation set where six summarize a metric from the manual coordinates, and six are selected from accuracy-coverage curves. The confidence threshold for determining whether to keep or discard a correction is 0.7, and road- and range-distance flagging thresholds are 0.10 and 1.0 km.

Sixteen thresholds encode physical quantities, external-data properties, or field-practice conventions. Varying each of the ten remaining user choices by ±50% changes flagging recall by at most 0.2% and usable-coordinate coverage by at most 1.5%, with unchanged correction accuracy (Table S2). For a new region, the calibration process re-measures the twelve calibrated thresholds from local expert coordinates. The ten insensitive defaults can be retained for future applications. Supplementary Methods S6 explains these choices.

### 2.4 Downstream-impact evaluation

We evaluate downstream consequences at two scales: the environment associated with each occurrence and the geographic range inferred for each species.

Environmental distance measures the error that a coordinate introduces into a species distribution model. Such a model uses predictor values extracted at an occurrence, not the coordinate itself. Positional error matters only when it changes those values (Gábor et al., 2019; Naimi et al., 2011). At each raw, GeoMend, and expert coordinate, we sampled the six continuous predictors commonly used in species distribution modeling: elevation, distance to stream, distance to primary forest, distance to secondary forest, and primary and secondary forest cover within 600 m. The land-cover predictors came from a 30 m annual reconstruction for Hispaniola from 1996 to 2022, read for the record’s collection year (Hong et al., 2025).

We combined the six predictor errors into one standardized environmental distance. For each predictor, we divided the difference from the expert value by the standard deviation across expert coordinates, then calculated the root mean square of the six standardized differences. Zero indicates that a coordinate occupies the same measured environment as the expert coordinate. We report this distance only for records GeoMend corrects.

The extent of occurrence (EOO) is the area of the minimum convex polygon around a species’ records and is used in IUCN Red List assessments (IUCN, 2012). A single grossly misplaced record can therefore inflate the apparent range. We calculated EOO from the original coordinates in the study dataset, GeoMend output, and expert coordinates, then compared the raw and corrected values with expert truth. This analysis included species with enough records to form a hull in all three sets: 149 orchids, 80 frogs, and 153 reptiles (Fig. 4). We also tested whether the raw or corrected coordinates placed the species in different categories of the 100, 5,000, and 20,000 km² thresholds associated with the Critically Endangered, Endangered, and Vulnerable categories under IUCN Criterion B1 (IUCN, 2012). This comparison is not a Red List assessment, which would also require evidence about threats, decline, fragmentation, and population, but it does provide an illustration of the potential consequences for conservation of correcting positional error. For every species that formed a hull under all three coordinate sets, we compared the raw and GeoMend threshold classes with the expert class and reported disagreement by taxon (Fig. 4b). Because newly georeferenced records and corrected coordinates both affect the result, we treat this analysis as secondary.

### 2.5 Implementation and reproducibility

GeoMend is an open-source Python command-line package under the MIT licence. Identical inputs and configuration produce identical outputs. The thresholds reside in one configuration file. Separate scripts reproduce calibration, sensitivity analyses, benchmarks, downstream evaluation, and figures. Source and data access are described in the peer-review statement.

Anthropic Claude Code v2.1.274 and OpenAI Codex GPT-6 were used during software development, documentation, analysis-script development, and manuscript editing. AI assistance was also used for drafting and refactoring the analysis and pipeline code and for writing tests, and for copy-editing the manuscript. All scientific content, prose, and analytical decisions belong to the authors. The authors reviewed all AI-assisted code and text, and the corresponding author takes responsibility for its accuracy, reproducibility, and intellectual content.

### 2.6 Benchmarking against existing tools

We ran GeoMend on the 5,000-locality Nature Plants benchmark without regional adjustment, comparing coverage and great-circle error against GPT-4o, GEOLocate, and county centroids (Rios & Bart, n.d.; Xie et al., 2025). On Hispaniola, GEOLocate received the locality, province, and country. We tested its road/highway option both enabled and disabled and cached all responses (Supplementary Methods S6).

## 3. Results

### 3.1 Evaluation using the Hispaniola records

Detection and correction are evaluated on held-out records (n = 6,322), including all orchids (n = 3,674), although the primary species-reference ranges use all expert coordinates (Section 2.2).

#### 3.1.1 Detection

GeoMend flags most genuinely misplaced coordinates with higher recall than a widely used flag-only baseline, CoordinateCleaner (Fig. 3a). Of the held-out coordinates more than 5 km from expert truth, it flags 92%, against 27% for CoordinateCleaner. Of those more than 1 km off, it flags 89% against 19%. Of the records that a human reviewer independently flagged, it recovers 80% against 10%. Unlike CoordinateCleaner, GeoMend also proposes a corrected coordinate. At the 5 km target recall is 94% for orchids, 92% for frogs, and 88% for reptiles, and at 1 km it is 94%, 89%, and 81%.

These are recall figures, which say how much of the real error each tool catches. Precision, the share of raised flags that are real errors, is a separate question, and on the greater-than-1-km target the two tools are close, 57% for GeoMend against 59% for CoordinateCleaner. GeoMend therefore catches three to five times as many real errors at essentially the same flag purity, and because a flagged record is kept with elevated uncertainty rather than discarded, a false flag widens a record’s error bound rather than removing the record.

The species-range detectors are assessed separately because they build each species’ reference range from expert coordinates and so are not fully independent (Section 2.2). We therefore reran detection under five reference sets, from the full expert table to no species reference at all (Table S5). Recall at the 5 km target spans 88% to 92% across all five. Rebuilding the reference from GeoMend’s own first-pass coordinates, which uses no expert data at all, holds recall at 92% and lowers endorsed coverage by 3.7 percentage points. Even with no species-level reference at all, recall is 88%, which is more than three times CoordinateCleaner’s 27%. This demonstrates that detection does not require expert coordinates.

#### 3.1.2 Correction

GeoMend’s corrections target a specific and common error mode. Field localities lie on roads and trails, but their coordinates are often digitized as a straight line drawn from a named place, which lands the point off the road and often far from the true collection location. GeoMend walks the stated distance along the road and offsets the result 50 m into adjacent habitat. Across Hispaniola, it accepted 1,212 corrections. Among accepted corrections with coordinates available for paired scoring, the median error falls from 2.19 km in the original coordinates to 0.82 km after GeoMend, and the fraction within 5 km rises from 88% to 90%. Of the 298 held-out corrections in the evaluation, 291 have complete coordinates for paired scoring, and seven lack a longitude. Among the complete pairs, the median error after correction is 0.98 km, 89% fall within 5 km of truth, and 68% improve. For the 197 improved records, the median error falls from 3.10 to 0.42 km. For the 94 that do not improve, the median increase in error is 1.83 km. The mechanism is visible in the N Pedernales series (Fig. 1b), where raw points fall on an artificial straight-north line and the road-derived coordinates follow the real curving road.

#### 3.1.3 Coverage and disposition

Each record is assigned to exactly one precision tier, from a trusted original at a median 0.70 km from truth to a country centroid near 93 km, according to the gazetteer used in this study, with the full per-tier breakdown in Table S3 (Fig. 2). The tiers give 74.8% of the records an endorsed coordinate, meaning an output in Tiers 0–3 that the pipeline recommends at its labelled precision. A further 25.0% are flagged for expert review, but retain their own coordinate, and 30 records (0.2%), which name no locatable place, receive none.

**Figure 2.**
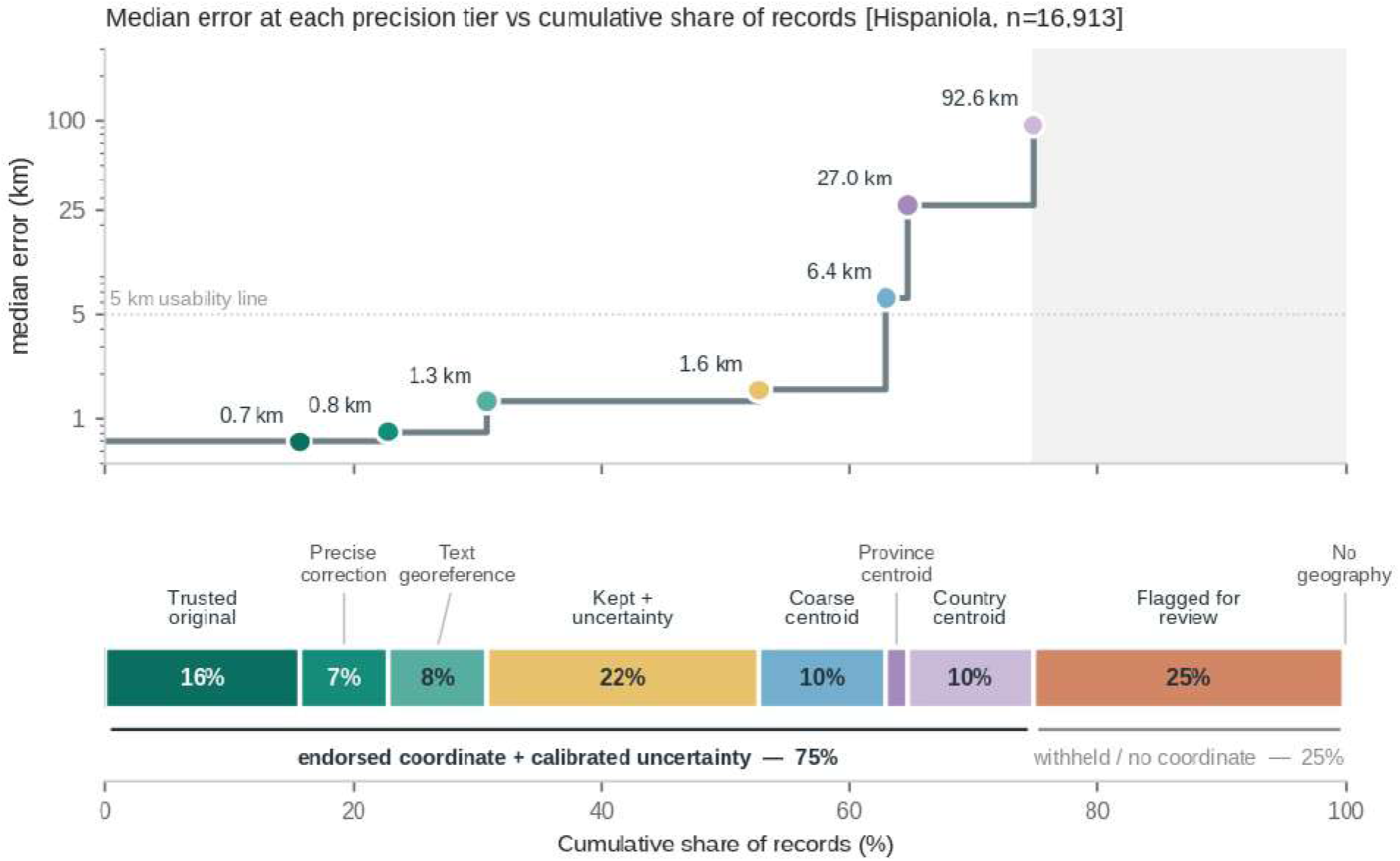
Coverage at precision on Hispaniola. The step plot relates each tier’s median error to cumulative coverage, and the aligned bar identifies the tier added at each step (n = 16,913).

Across tiers, the calibrated radius contains the expert coordinate at close to its stated rate: 90% for kept-with-uncertainty and 93% for the coarse gazetteer tier on the full validation set, and 84 to 95% for the administrative fallback (Table S3). Source-reported uncertainty, if provided, is retained separately. For an unchanged coordinate, GeoMend reports the larger of the supplied and the modelled radius, whereas a moved point receives only its newly calibrated radius.

What is leftover limits precise placement rather than coverage. The records that reach the coarse tiers or stay flagged are dominated by bare localities, which name a place and give no distance or bearing, so there is nothing to resolve to a point. Adding further global gazetteers did not improve coverage in our analysis: relative to GeoNames, OpenStreetMap place names resolve a further 1.7% of these bare localities, and the NGA GNS gazetteer only 0.3%.

#### 3.1.4 Downstream range recovery

Correction moves records toward the environments the experts assigned them (Fig. 4a). Across the 1,173 corrected records, the median standardized environmental distance to the expert coordinate falls from 0.436 to 0.207, a reduction of 53% (95% CI for the median difference, 0.201 to 0.252). The error falls on each of the six predictors. The largest relative change is in elevation, whose median absolute error falls from 63 to 18 m, a reduction of 72%. The distance to primary forest follows, falling from 490 to 210 m, then the distance to the stream, from 416 to 203 m. Among 324 species with at least five records, 124 move closer to the expert environment, 22 move further, and 178 remain unchanged (sign test, p = 1.9 × 10^-18^). The gain is clearest for reptiles, whose EOO ratio barely moves, from 1.02 to 1.01, while their median environmental distance falls from 0.43 to 0.22 across 857 corrected records. Environmental distance asks whether the correction recovers the predictor values sampled at the expert coordinates.

Correcting the raw coordinates also moves the extent of occurrence toward expert truth for every taxon, without inflating ranges that are already well defined (Fig. 4a). Against expert hulls built from every expert-georeferenced record, orchid EOO moves from 8.36 times the truth to 1.98 times, frog EOO from 2.01 to 0.96 times using corrections alone, and to 1.12 times once text-georeferenced points are added, and reptile EOO from 1.02 to 1.01 times. Because expert hulls contain some records that GeoMend abstains from placing, we also matched their record support exactly: the corresponding raw-toGeoMend ratios are 12.24 to 2.89 for orchids, 2.28 to 1.09 for frogs, and 1.17 to 1.16 for reptiles. The conclusion is unchanged, although the absolute ratios depend on which records support the hull. We also tested per species and found that the raw frog ranges scatter far off the one-to-one line with truth, while the corrected ranges are more closely aligned (Fig. S1).

GeoMend also enables a new range estimate for species that the original coordinates cannot assess. Seventeen species gain a convex hull and, with it, an extent of occurrence and a better basis for habitat estimation. The expert makes sixty-five more species modelable that GeoMend leaves flagged, declining to place them on weak evidence, and we return to that gap in the Discussion.

Correction shifts the EOO class associated with IUCN Criterion B1 in the same direction (Section 2.4). On the 382 species that form a hull under all three coordinate sets, the class based on original coordinates differs from the expert-truth class for 26% of frogs, 35% of orchids, and 26% of reptiles, and GeoMend reduces this to 16%, 26%, and 23%. In the orchid *Epidendrum miserrimum*, a single outlier inflates EOO from 2,125 km² to 871,216 km², moving it from the Endangered-associated threshold to above the threatened thresholds. GeoMend corrects this outlier, aligning conservation status with the expert (Fig. 4b).

#### 3.1.5 Comparison with the standard georeferencing service

GeoMend places Hispaniola records more accurately than the standard automated georeferencing service (Section 2.6). On the 568 records where GeoMend, GEOLocate, and the original coordinate all resolve, GeoMend’s coordinate falls a median of 0.67 km from truth against 10.69 km for GEOLocate and 2.07 km for the raw coordinate, and places 56% of those records within 1 km against 5% for GEOLocate. Across the records that GEOLocate itself resolves, the error distribution has a heavy tail, with a median of 7.0 km and a mean of 36 km. The pattern of those errors, which lie along the stated bearing at roughly the stated distance, is a demonstration of a straight-line placement rather than a road-following placement. Running GEOLocate with its road and highway correction enabled produced identical results. GEOLocate returned no usable uncertainty radius for any of these records.

### 3.2 Evaluation using the Nature Plants benchmark

GeoMend’s georeferencing generalizes beyond Hispaniola. Its most common error on the benchmark is 4.25 km, against 9.00 km for GPT-4o and 9.82 km for GEOLocate (Fig. 3b). Run on the Nature Plants benchmark with the same defaults and no region-specific adjustment, its coordinates are closer to the published coordinates than GPT-4o’s for two-thirds of the records that both tools place (67%, 378 of 566), with a median error of 4.44 km against 10.76 km for GPT-4o, 11.97 km for GEOLocate, and 26.4 km for the county-centroid baseline.

**Figure 3.**
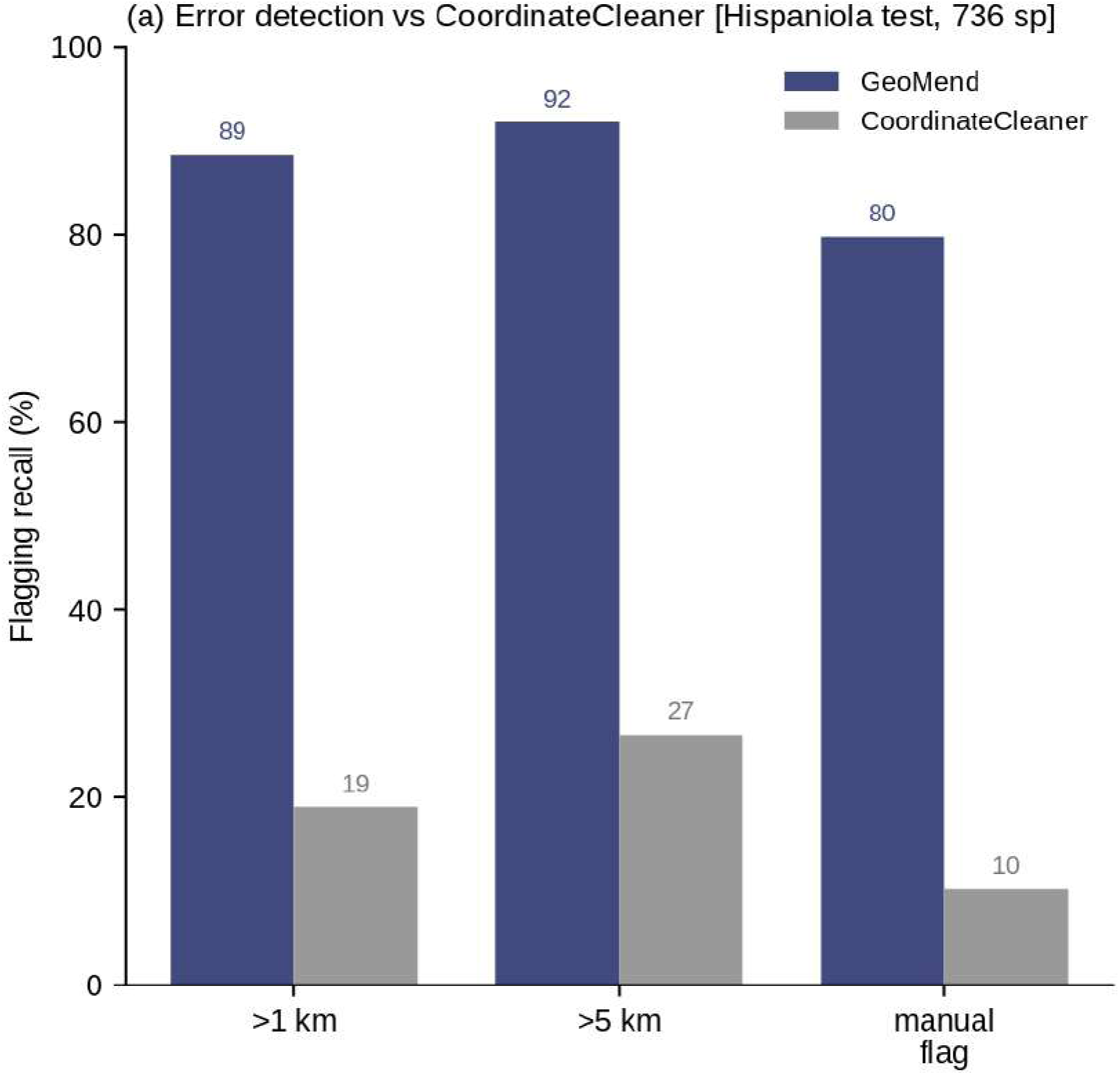
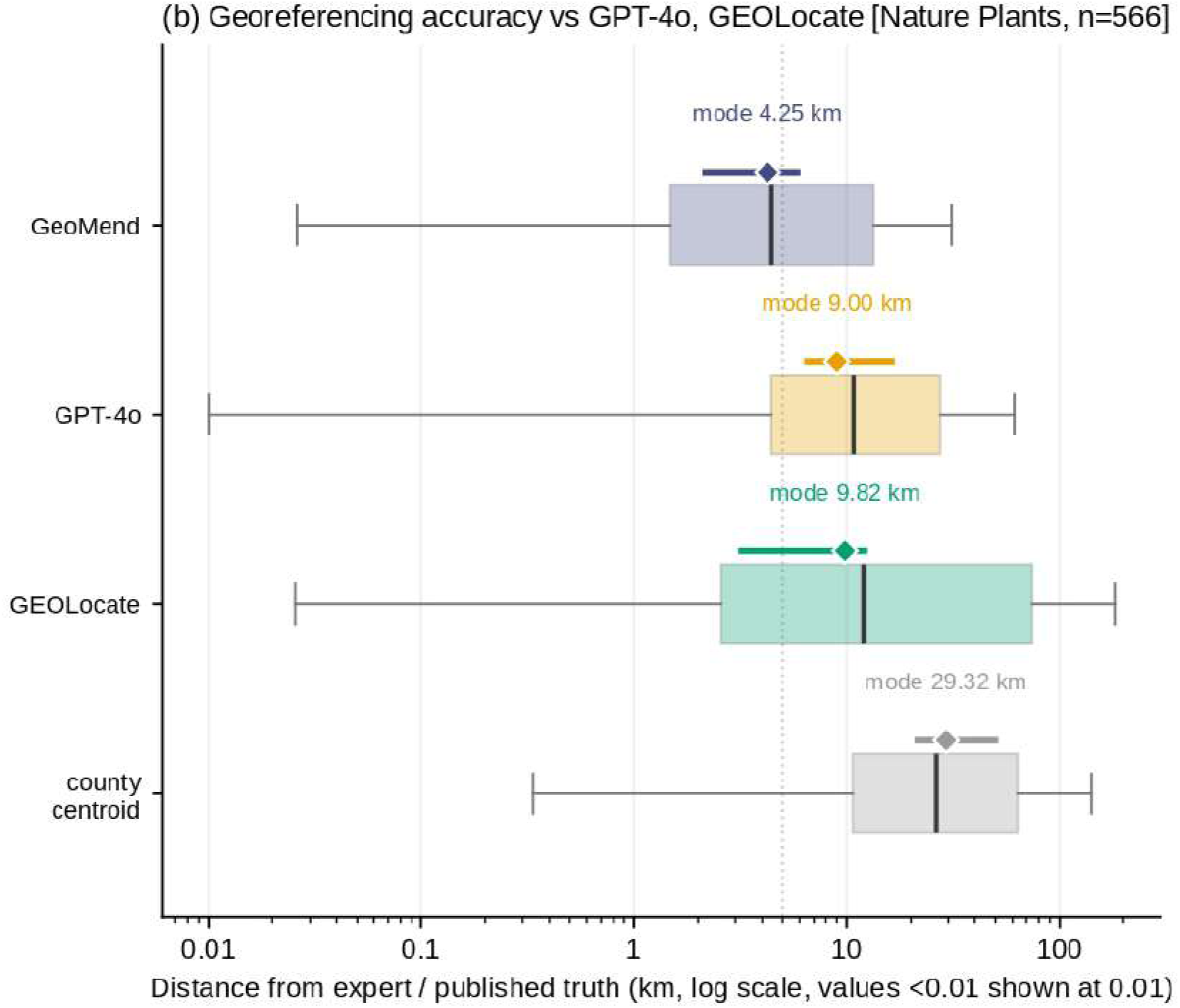
Comparison with existing tools. **(a)** Detection recall versus CoordinateCleaner on held-out Hispaniola records under three matched error definitions. **(b)** Error on the 566 Nature Plants records placed by all four georeferencers. Boxes show quartiles, whiskers use a logarithmic axis, and diamonds show bootstrapped modal error. For display on the logarithmic axis, errors below 0.01 km are plotted at 0.01 km, and all statistics use the original values.

**Figure 4.**
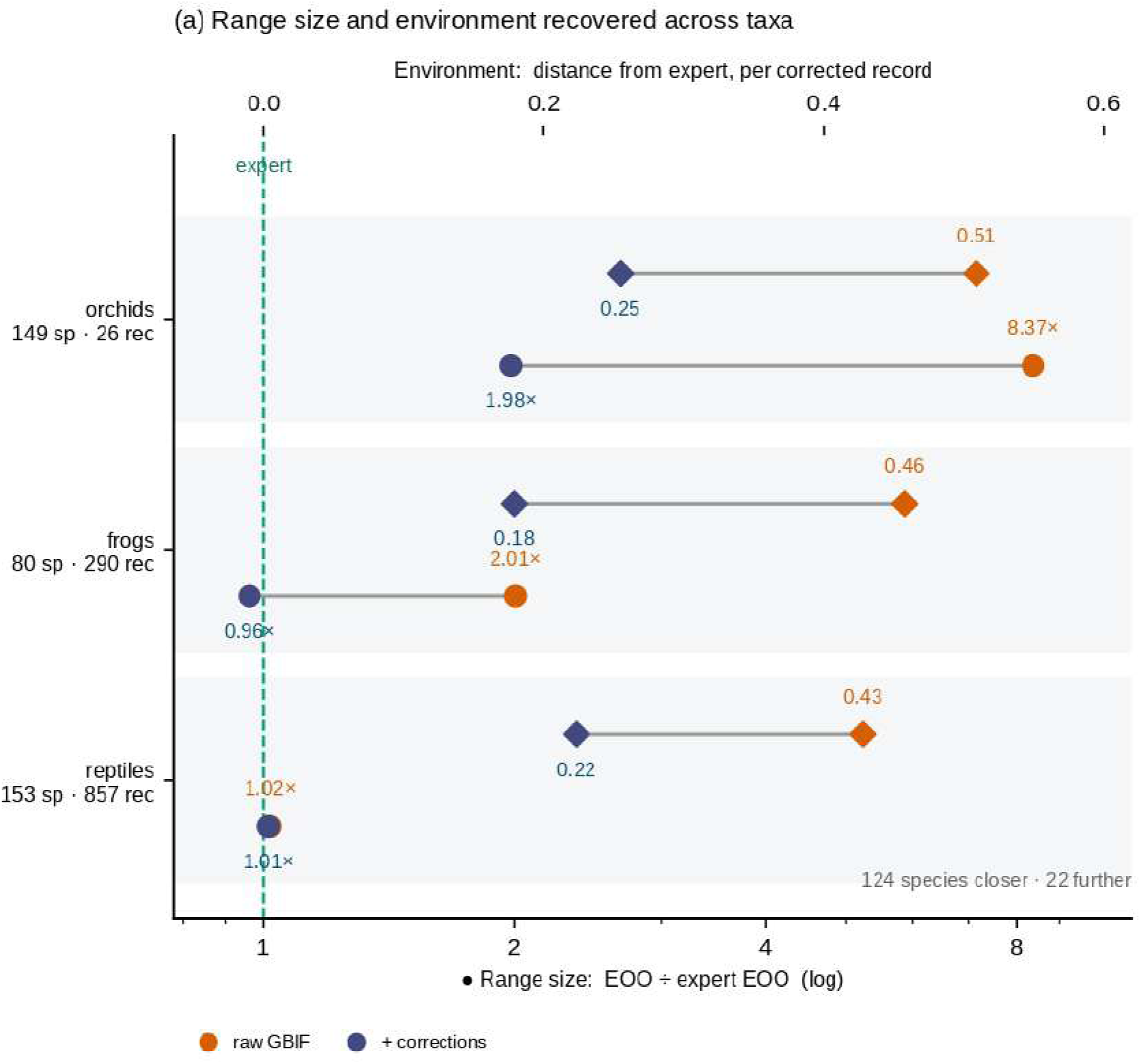
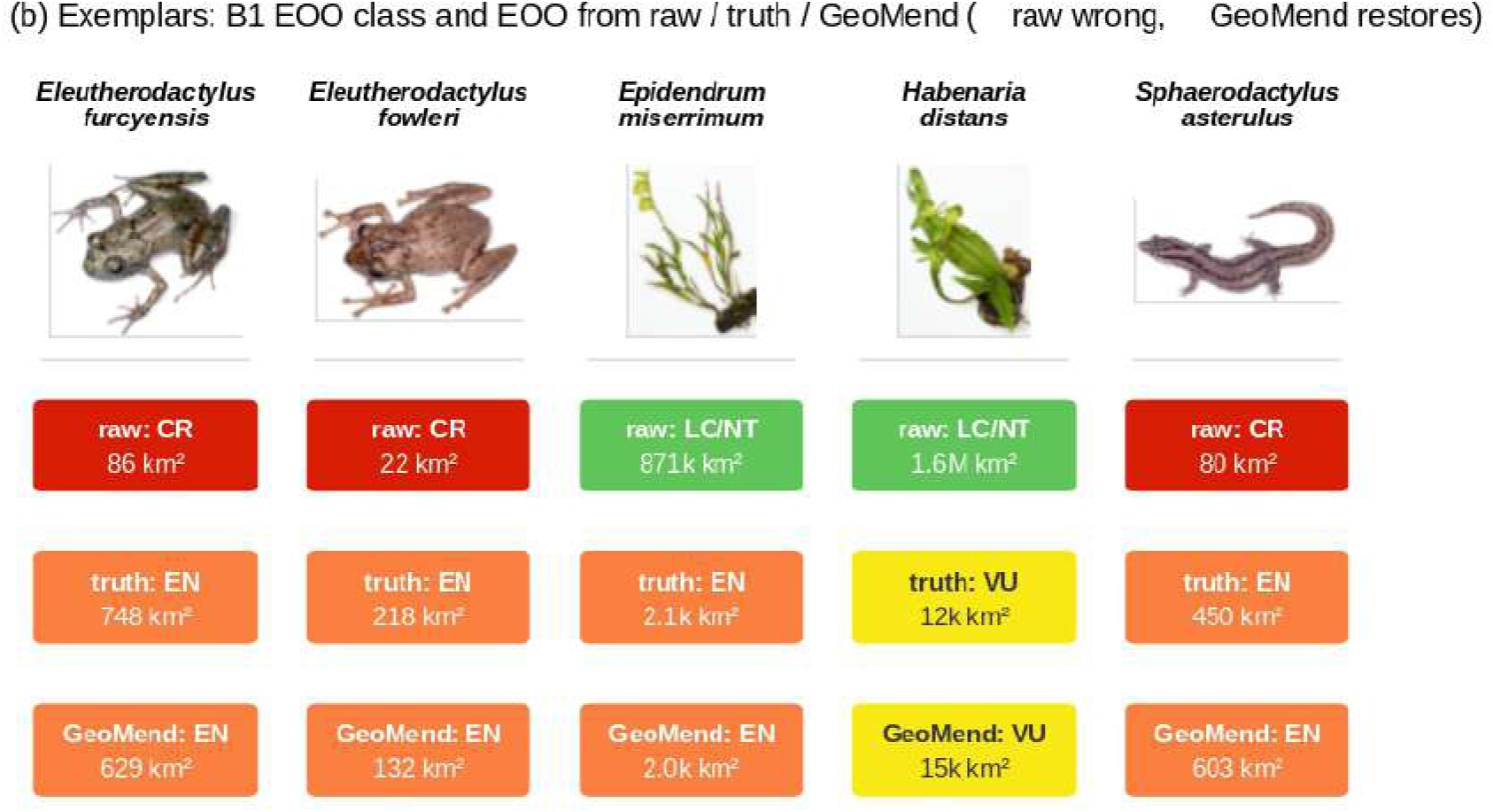
Downstream range recovery and the environment of corrected records. **(a)** How far GeoMend moves occurrence data toward expert truth, by taxon, on two measures. Diamonds, top axis, give the distance from the expert coordinate in environmental space for the records GeoMend corrects, where zero means the record occupies the environment of the expert coordinate. Circles, bottom axis, give the aggregate extent of occurrence relative to expert truth on a logarithmic scale. Both axes are aligned, so the dashed line marks the expert. Environmental distance and the six predictors behind it are defined in Section 2.4. Range-size ratios including text-georeferenced points are 2.04 for orchids, 1.12 for frogs, and 1.06 for reptiles. **(b)** Five species whose EOO class associated with IUCN Criterion B1 changes, comparing raw, expert, and GeoMend ranges. Photographs: *Epidendrum miserrimum*, Father Alejandro Sánchez†. *Habenaria distans*, Ernesto Mújica. *Eleutherodactylus furcyensis*, *Eleutherodactylus fowleri*, and *Sphaerodactylus asterulus*, S. Blair Hedges. †Deceased.

GeoMend places 69% of benchmark localities, or 11% when restricted to precise tiers, with tier labels allowing users to trade coverage for precision. Unlike the language model, every coarse placement carries an explicit uncertainty radius calibrated on Hispaniola. These require independent calibration elsewhere. All 72 countries in this analysis used the same defaults.

### 3.3 Species without georeferenced records

GeoMend georeferences records lacking coordinates, making them available for spatial analysis. These recovered records fall where data are scarcest. We demonstrate this first on Hispaniola, then across the global backlog. On Hispaniola it recovers 3,086 records that had no coordinate, and 68 species with no usable original coordinate in the study dataset gain a mapped occurrence, among them 27 frogs, 34 orchids, and 7 reptiles. A separate set of 17 species gains an extent of occurrence in Section 3.1.4. These sets are not nested because they measure different starting conditions: the 68 begin with no mapped occurrence, whereas the 17 begin with fewer than the three points needed to form a convex hull. The recovered records concentrate on rare species, with about 30% of recovered orchid records going to species represented by fewer than five original coordinates in the study dataset.

The same data gap exists globally. Of 1,491,014 species with preserved specimens in the full snapshot, 410,649 (27.5%) carry no coordinate on any record. At least one record has locality text GeoMend can read for 262,794 of these coordinate-less species, making that subset potentially addressable by species distribution models and IUCN assessment.

GeoMend returns a first coordinate for nearly one-third of previously unaddressable species. Applied to all 898,176 records, it places 242,033 records from 82,180 species (31%) across 233 of 245 countries (Fig. 5a). Every one of those 233 countries gains species at the country or regional grain. At the finer 100 km grain, precise placements increase richness in 2,931 grid cells, including 219 cells that each gain at least ten previously invisible species. These gains concentrate in the megadiverse tropics rather than in already well-collected regions.

**Figure 5.**
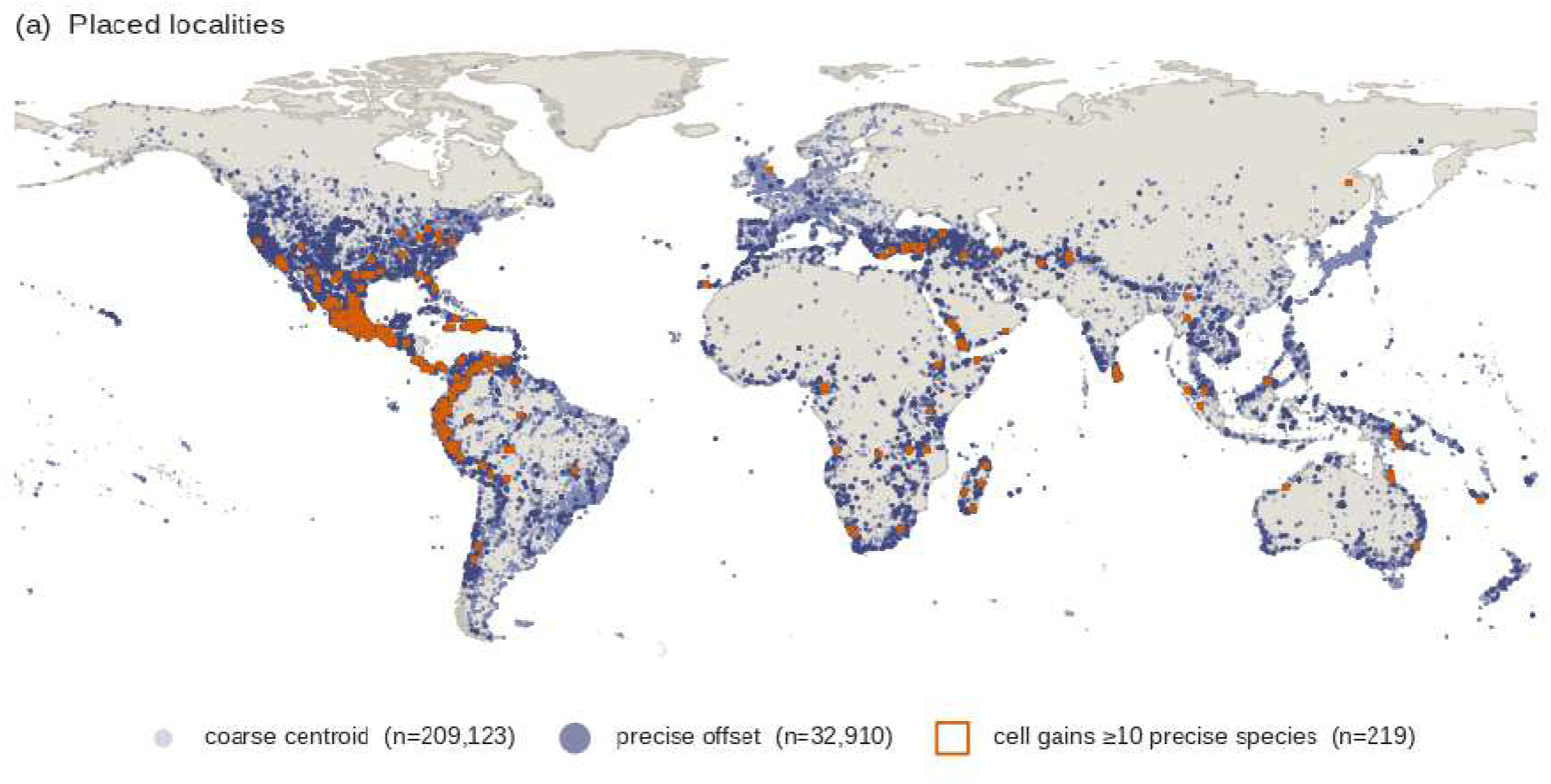
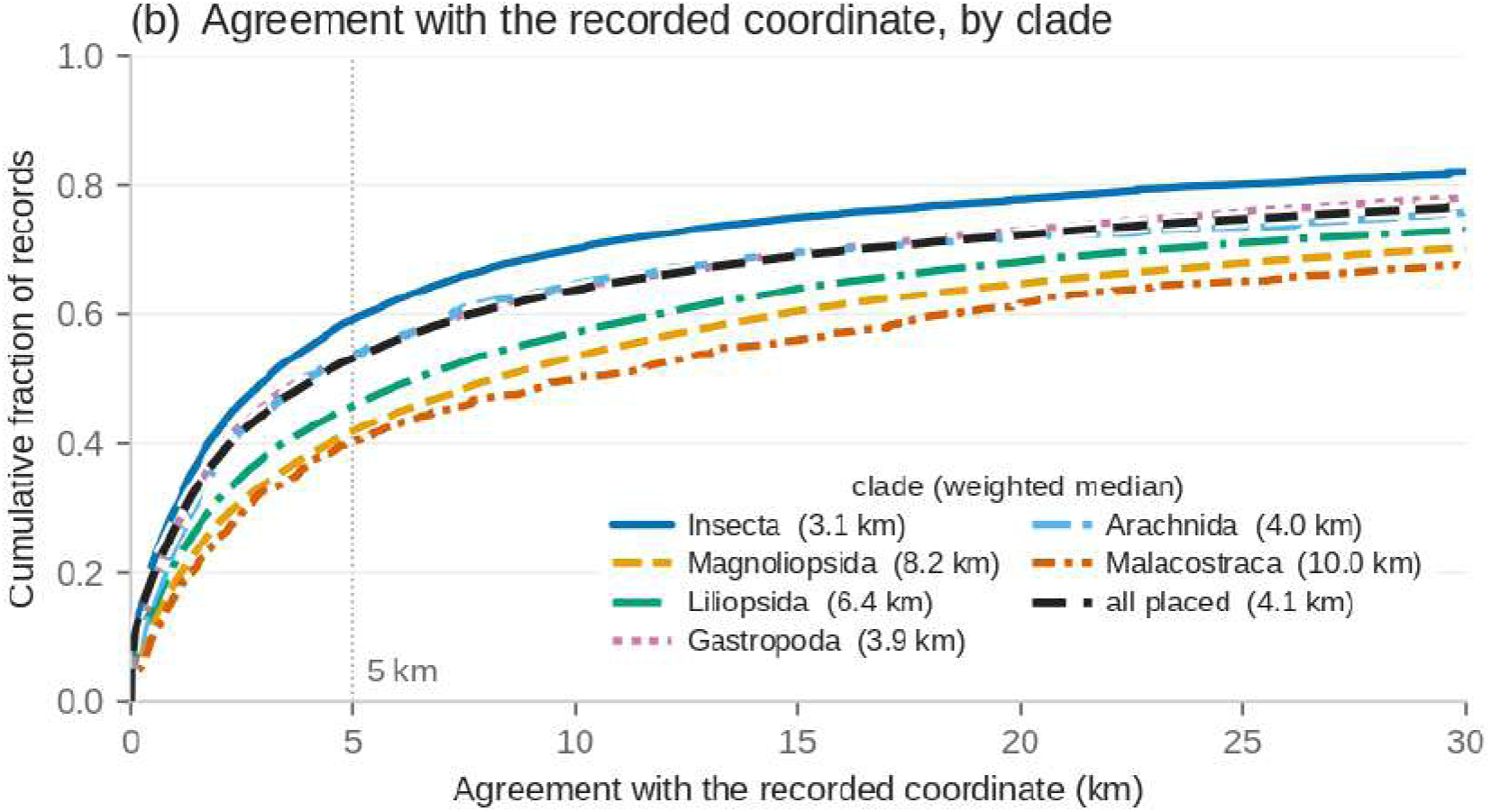
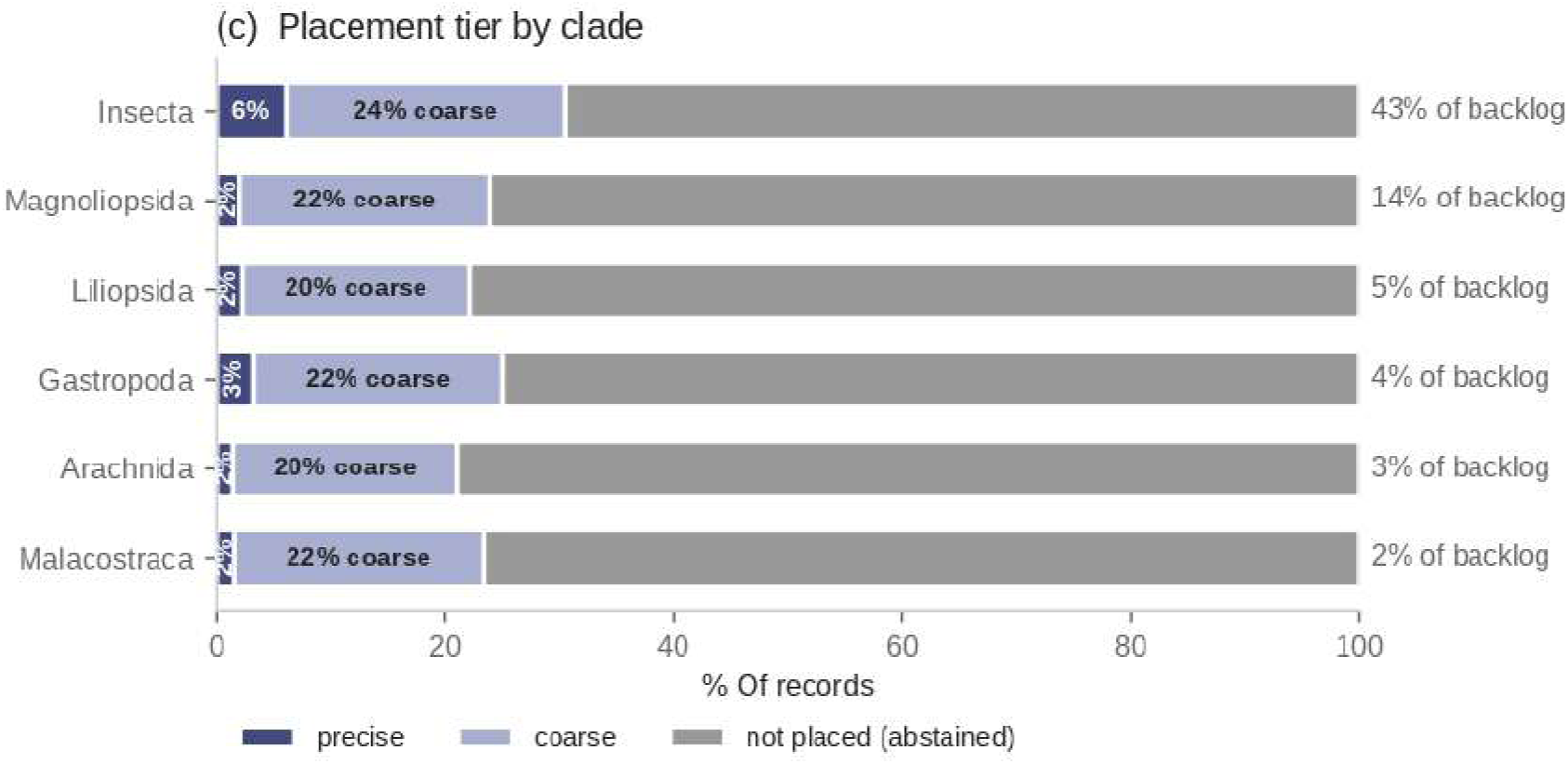
Global recovery from the pinned GBIF snapshot. **(a)** Placed localities by precision tier. **(b)** Post-stratified agreement with recorded donor coordinates by clade. This is agreement with similar coordinates, not independent accuracy of the global backlog placement. **(c)** Placement tier by clade, showing the invertebrate- and plant-dominated backlog.

The global backlog of records lacking coordinates is concentrated in the taxa that global analyses most often overlook. Insects alone are 43% of it, and flowering plants a further 15%, while the squamates, amphibians, birds, and mammals that dominate expert range maps are together under 4% (Fig. 5c). The species GeoMend returns are therefore disproportionately invertebrates and plants for which no expert-drawn range exists.

Where a placed record from the donor set can be checked against the coordinate it already carries, GeoMend’s text-derived position agrees with a weighted median of 4.1 km, and with 2.6 km on the precise tier, with 53% of records within 5 km (Fig. 5b). This is a separate test to show agreement, not independent accuracy, because pre-1993 coordinates were frequently desk-georeferenced from the same locality string GeoMend reads, so the Hispaniola records of Section 3.1 remain independent evidence (Xie et al., 2025). GeoMend declines far more than it places, leaving 73% of these backlog records unplaced and giving a precise coordinate to only about 3% of species, and a further 12% of records name no country at all and so are currently unplaceable by GeoMend.

## 4. Discussion

GeoMend’s principal contribution is to make previously ungeoreferenced occurrence records available for quantitative analysis with explicitly stated precision. It provided first coordinates for 82,180 species across 233 countries and, on Hispaniola, improved positional accuracy and made EOO estimable for 17 previously unassessable species. The gain is therefore not only accuracy but also access to occurrence information otherwise excluded from spatial biodiversity analyses (Hortal et al., 2015; Meyer et al., 2015).

GeoMend formalizes the decisions that expert georeferencers make repeatedly: whether locality evidence supports a correction, what spatial precision that evidence warrants, and when a record should remain unresolved. These are the decisions georeferencing best practice already asks a curator to document (Chapman & Wieczorek, 2020; Wieczorek et al., 2004), and encoding them in explicit rules makes their application consistent, reproducible, and quick to compute across large datasets. It does not remove the need for expert judgment. The precision tiers instead separate records that stated rules can treat consistently from those that require individual review, because a coordinate carrying no measure of precision can imply more geographic knowledge than the locality description supports.

GeoMend corrects positional error, not the geographic bias of collecting. Records still concentrate near roads and accessible places because those are the places collectors visited (Daru et al., 2018; He et al., 2025; Hughes et al., 2021; Monsarrat et al., 2018; Ruete, 2015). Road-derived coordinates therefore require particular care in distribution modelling (Baker et al., 2024; Kadmon et al., 2004). Among the 1,352 records receiving such coordinates, the 50 m habitat offset reduced but did not eliminate differences in composition measured with the annual land-cover reconstruction (Hong et al., 2025).

Natural-history information offers a direct route to improving these decisions, such as off-road habitat offset or precise placement. More detailed information on habitat association, elevation, dispersal, and collecting practice could further constrain candidate locations. For example, a road-derived coordinate could be placed in adjacent habitat appropriate to the taxon rather than at a fixed distance from the road. Such extensions would retain the reproducibility of explicit rules while incorporating biological differences among species (Chapman & Wieczorek, 2020; Stenhouse et al., 2025).

Weak locality evidence limits how precisely any automated method can place a record (Chapman & Wieczorek, 2020; Wieczorek et al., 2004). Bare or descriptive locality text may remain unresolved without historical maps or regional knowledge, although improved name matching, map interpretation, and parsing may recover some records (GeoNames, n.d.; Wijegunarathna et al., 2025). Precision tiers must also match the scale of analysis because the effect of locational uncertainty depends on predictor resolution and spatial heterogeneity (Gábor et al., 2022; Mitchell et al., 2017). Tiers 0, 1a, and 1b support finescale uses, whereas administrative centroids support only coarser summaries (Gábor et al., 2022; Graham et al., 2008).

The uncertainty radii are calibrated on Hispaniola and require independent calibration elsewhere (Wieczorek et al., 2004; Xie et al., 2025). Detection does not depend on expert coordinates (Table S5), but the radii do. Calibrating them globally does not require expert coordinates in every region. Gazetteers use the same place types everywhere, and the donor comparison of Section 3.3 estimates the same error distributions from coordinates that the records already carry, as agreement rather than accuracy. Whether radii are better pooled across regions or stratified by region and taxon remains open (Stenhouse et al., 2025). Until they are calibrated locally, abstention protects the user.

GeoMend is a first step toward clearing that backlog at scale. It placed 898,176 records in about one minute on 16 processor cores and carries no extra cost per record, whereas language-model georeferencing costs about $0.001 per record (Xie et al., 2025) and manual georeferencing takes minutes to hours (Stenhouse et al., 2025).

Altogether, resolving the remaining backlog will require including more information from improved historical maps, natural history information, and experimentation with stratification (Chapman & Wieczorek, 2020; Stenhouse et al., 2025). Large language models could coordinate verifiable components: interpreting locality descriptions, retrieving maps, applying deterministic procedures, and referring unresolved cases to regional experts (Wijegunarathna et al., 2025). Calibrated uncertainty and abstention remain essential because model confidence alone does not establish support for a placement (Kirchhof et al., 2025). Collections hold 1.2 to 2.1 billion specimens (Ariño, 2010; Johnson et al., 2023), most of them neither digitized nor mapped. Every one of them is a record of a species in a habitat, and extinction risk cannot be judged for a species whose habitats are unknown. Mapping the backlog would bring these unmapped species into focus for the first time and allow for previously unattainable extinction-risk assessment (Hortal et al., 2015; IUCN, 2012).

## Notes

https://osf.io/mcdhb/?view_only=ab3de27644ed437994d1cf3c30da079b

### Competing Interest Statement

The authors have declared no competing interest.

### Funding

- U.S. National Science Foundation: 2326013, 2326014

## References

Aldana-Bobadilla, E., Molina-Villegas, A., Lopez-Arevalo, I., Reyes-Palacios, S., Muñiz-Sanchez, V., & Arreola-Trapala, J. (2020). Adaptive geoparsing method for toponym recognition and resolution in unstructured text. Remote Sensing, 12(18), 3041. 10.3390/rs12183041

Ariño, A. H. (2010). Approaches to estimating the universe of natural history collections data. Biodiversity Informatics, 7(2), 81–92. 10.17161/bi.v7i2.3991

Baker, D. J., Maclean, I. M. D., & Gaston, K. J. (2024). Effective strategies for correcting spatial sampling bias in species distribution models without independent test data. Diversity and Distributions, 30(3), e13802. 10.1111/ddi.13802

Barber, A., Lafferty, D., & Landrum, L. R. (2013). The SALIX method: A semiautomated workflow for herbarium specimen digitization. TAXON, 62(3), 581–590. 10.12705/623.16

Blagoderov, V., Kitching, I., Livermore, L., Simonsen, T., & Smith, V. (2012). No specimen left behind: Industrial scale digitization of natural history collections. ZooKeys, 209, 133–146. 10.3897/zookeys.209.3178

Center for Biodiversity, Temple University. (2025). CBTU. https://biodiversitycenter.org/

Chapman, A. D., & Wieczorek, J. R. (2020). Georeferencing best practices. GBIF Secretariat. 10.15468/doc-gg7h-s853

Daru, B. H., Park, D. S., Primack, R. B., Willis, C. G., Barrington, D. S., Whitfeld, T. J. S., Seidler, T. G., Sweeney, P. W., Foster, D. R., Ellison, A. M., & Davis, C. C. (2018). Widespread sampling biases in herbaria revealed from large-scale digitization. New Phytologist, 217(2), 939–955. 10.1111/nph.14855

Elith, J., Phillips, S. J., Hastie, T., Dudík, M., Chee, Y. E., & Yates, C. J. (2011). A statistical explanation of MaxEnt for ecologists. Diversity and Distributions, 17(1), 43–57. 10.1111/j.1472-4642.2010.00725.x

Fernando, A., Ranathunga, S., Stock, K., Prasanna, R., & Jones, C. B. (2026). Georeferencing complex relative locality descriptions with large language models. International Journal of Geographical Information Science, 1–33. 10.1080/13658816.2026.2613291

Gábor, L., Jetz, W., Lu, M., Rocchini, D., Cord, A., Malavasi, M., Zarzo-Arias, A., Barták, V., & Moudrý, V. (2022). Positional errors in species distribution modelling are not overcome by the coarser grains of analysis. Methods in Ecology and Evolution, 13(10), 2289–2302. 10.1111/2041-210X.13956

Gábor, L., Moudrý, V., Lecours, V., Malavasi, M., Barták, V., Fogl, M., Šímová, P., Rocchini, D., & Václavík, T. (2019). The effect of positional error on fine scale species distribution models increases for specialist species. Ecography, 43(2), 256–269. 10.1111/ecog.04687

GBIF.org. (2026a). GBIF Occurrence Download: Occurrence snapshot, 1 July 2026. 10.15468/dl.teccpa

GBIF.org. (2026b). GBIF Occurrence Download: Occurrence snapshot, 1 August 2026. 10.15468/dl.fs3btq

Geofabrik GmbH. (n.d.). Geofabrik download server: Haiti and dominican republic extract. https://download.geofabrik.de/central-america/haiti-and-domrep.html.

GeoNames. (n.d.). GeoNames geographical database. https://www.geonames.org.

Graham, C. H., Elith, J., Hijmans, R. J., Guisan, A., Peterson, A. T., Loiselle, B. A., & The NCEAS Predicting Species Distributions Working Group. (2008). The influence of spatial errors in species occurrence data used in distribution models. Journal of Applied Ecology, 45(1), 239–247. 10.1111/j.1365-2664.2007.01408.x

Guisan, A., Tingley, R., Baumgartner, J. B., Naujokaitis-Lewis, I., Sutcliffe, P. R., Tulloch, A. I. T., Regan, T. J., Brotons, L., McDonald-Madden, E., Mantyka-Pringle, C., Martin, T. G., Rhodes, J. R., Maggini, R., Setterfield, S. A., Elith, J., Schwartz, M. W., Wintle, B. A., Broennimann, O., Austin, M., … Buckley, Y. M. (2013). Predicting species distributions for conservation decisions. Ecology Letters, 16(12), 1424–1435. 10.1111/ele.12189

Halterman, A. (2017). Mordecai: Full text geoparsing and event geocoding. The Journal of Open Source Software, *2*(9), 91. 10.21105/joss.00091

He, J., Yang, W., You, Q., Hu, Q., Cong, M., Tian, C., & Ma, K. (2025). Impacts of road networks on the geography of floristic collections in China. Plant Diversity, 47(3), 403–414. 10.1016/j.pld.2025.02.001

Hedges, S. B. (2013). Caribmap: A mobile-friendly web resource for caribbean maps that facilitates quantitative historical analysis. Journal of Map and Geography Libraries, 9, 350–360. 10.1080/15420353.2013.804476

Hong, F., Hedges, S. B., Yang, Z., Suh, J. W., Qiu, S., Timyan, J., & Zhu, Z. (2025). Decoding primary forest changes in Haiti and the Dominican Republic using Landsat time series. Remote Sensing of Environment, 318, 114590. 10.1016/j.rse.2024.114590

Hortal, J., Bello, F. de, Diniz-Filho, J. A. F., Lewinsohn, T. M., Lobo, J. M., & Ladle, R. J. (2015). Seven shortfalls that beset large-scale knowledge of biodiversity. Annual Review of Ecology, Evolution, and Systematics, 46, 523–549. 10.1146/annurev-ecolsys-112414-054400

Hughes, A. C., Orr, M. C., Ma, K., Costello, M. J., Waller, J., Provoost, P., Yang, Q., Zhu, C., & Qiao, H. (2021). Sampling biases shape our view of the natural world. Ecography, 44, 1259–1269. 10.1111/ecog.05926

IUCN. (2012). IUCN red list categories and criteria: Version 3.1 (Second). IUCN.

Jin, J., & Yang, J. (2020). BDcleaner: A workflow for cleaning taxonomic and geographic errors in occurrence data archived in biodiversity databases. Global Ecology and Conservation, 21, e00852. 10.1016/j.gecco.2019.e00852

Johnson, K. R., Owens, I. F. P., et al. (2023). A global approach for natural history museum collections. Science, 379(6638), 1192–1194. 10.1126/science.adf6434

Kadmon, R., Farber, O., & Danin, A. (2004). Effect of roadside bias on the accuracy of predictive maps produced by bioclimatic models. Ecological Applications, 14(2), 401–413. 10.1890/02-5364

Kirchhof, M., Kasneci, G., & Kasneci, E. (2025). Position: Uncertainty quantification needs reassessment for large-language model agents. Proceedings of the 42nd International Conference on Machine Learning (ICML), Proceedings of Machine Learning Research, 267, 81665–81677. https://proceedings.mlr.press/v267/kirchhof25b.html

Margules, C. R., & Pressey, R. L. (2000). Systematic conservation planning. Nature, 405, 243–253. 10.1038/35012251

Meyer, C., Kreft, H., Guralnick, R., & Jetz, W. (2015). Global priorities for an effective information basis of biodiversity distributions. Nature Communications, 6, 8221. 10.1038/ncomms9221

Mitchell, P. J., Monk, J., & Laurenson, L. (2017). Sensitivity of fine-scale species distribution models to locational uncertainty in occurrence data across multiple sample sizes. Methods in Ecology and Evolution, 8(1), 12– 21. 10.1111/2041-210X.12645

Monsarrat, S., Boshoff, A. F., & Kerley, G. I. H. (2018). Accessibility maps as a tool to predict sampling bias in historical biodiversity occurrence records. Ecography, 42(1), 125–136. 10.1111/ecog.03944

Naimi, B., Skidmore, A. K., Groen, T. A., & Hamm, N. A. S. (2011). Spatial autocorrelation in predictors reduces the impact of positional uncertainty in occurrence data on species distribution modelling. Journal of Biogeography, 38(8), 1497–1509. 10.1111/j.1365-2699.2011.02523.x

National Geospatial-Intelligence Agency, & U.S. Board on Geographic Names. (n.d.). NGA GEOnet names server (GNS). https://geonames.nga.mil.

OpenStreetMap contributors. (n.d.). OpenStreetMap. https://www.openstreetmap.org.

Patten, N. N., Gaynor, M. L., Soltis, D. E., & Soltis, P. S. (2024). Geographic and taxonomic occurrence R-based scrubbing (gatoRs): An R package and workflow for processing biodiversity data. Applications in Plant Sciences, 12(2), e11575. 10.1002/aps3.11575

Ravenscroft, H., Gardner, J., Perktas, U., & Saupe, E. E. (2025). Release the HOGS: An unsupervised marker extraction, classification and georeferencing approach for biodiversity data. Methods in Ecology and Evolution, 16(11), 2552–2559. 10.1111/2041-210X.70166

Ribeiro, B. R., Velazco, S. J. E., Guidoni-Martins, K., Tessarolo, G., Jardim, L., Bachman, S. P., & Loyola, R. (2022). bdc: A toolkit for standardizing, integrating and cleaning biodiversity data. Methods in Ecology and Evolution, 13(7), 1421–1428. 10.1111/2041-210X.13868

Rios, N. E., & Bart, H. L. (n.d.). GEOLocate: A platform for georeferencing natural history collections data. Tulane University Biodiversity Research Institute; https://www.geo-locate.org.

Ruete, A. (2015). Displaying bias in sampling effort of data accessed from biodiversity databases using ignorance maps. Biodiversity Data Journal, 3. 10.3897/bdj.3.e5361

Shirey, V., Seppälä, S., Branco, V., & Cardoso, P. (2019). Current GBIF occurrence data demonstrates both promise and limitations for potential red listing of spiders. Biodiversity Data Journal, 7. 10.3897/bdj.7.e47369

Stenhouse, A., Fisher, N., Lepschi, B., Schmidt-Lebuhn, A., Rodriguez, J., Turco, F., Reeson, A., Paris, C., & Thrall, P. H. (2025). A vision of human– AI collaboration for enhanced biological collection curation and research. BioScience, 75(6), 457–471. 10.1093/biosci/biaf021

Vollmar, A., Macklin, J. A., & Ford, L. (2010). Natural history specimen digitization: Challenges and concerns. Biodiversity Informatics, 7(2). 10.17161/bi.v7i2.3992

Wieczorek, J., Guo, Q., & Hijmans, R. J. (2004). The point-radius method for georeferencing locality descriptions and calculating associated uncertainty. International Journal of Geographical Information Science, 18(8), 745–767. 10.1080/13658810412331280211

Wijegunarathna, K., Stock, K., & Jones, C. B. (2025). Large multi-modal model cartographic map comprehension for textual locality georeferencing. 13th International Conference on Geographic Information Science (GIScience 2025), Leibniz International Proceedings in Informatics (LIPIcs), 346. 10.4230/LIPIcs.GIScience.2025.12

Xie, Y., Park, D. S., Sinnott-Armstrong, M. A., Ho, J., Chen, T., Weakley, A. S., Aguirre Lopez, L. J., Choi, J., Laitinen, M. M., Steeves, N. A., Huang, C. H., Xu, R., & Feng, X. (2025). Using large language models to address the bottleneck of georeferencing natural history collections. Nature Plants, 11(12), 2446–2450. 10.1038/s41477-025-02162-y

Zizka, A., Silvestro, D., Andermann, T., Azevedo, J., Duarte Ritter, C., Edler, D., Farooq, H., Herdean, A., Ariza, M., Scharn, R., Svantesson, S., Wengström, N., Zizka, V., & Antonelli, A. (2019). CoordinateCleaner: Standardized cleaning of occurrence records from biological collection databases. Methods in Ecology and Evolution, 10(5), 744–751. 10.1111/2041-210X.13152

