## Supplemental Information for "Correcting and georeferencing natural history occurrence records with calibrated uncertainty"

### Supplementary Methods

#### S1. Detector thresholds

A record with a coordinate is flagged as suspect if any of four criteria is met.

1. Physical plausibility. Flags a coordinate that falls in a water body such as the ocean, that sits more than 0.10 km from the nearest road, or whose elevation disagrees with the collector-recorded elevation by more than 150 m.

2. Range consistency. Using a trusted reference set of coordinates for each species, flags a coordinate more than 1 km outside the species' range polygon, one whose distance to the range centroid exceeds 0.5 times that species' own 95% spread, or one whose distance to its third-nearest same-species record exceeds that species' own 95th-percentile third-nearest-neighbour distance, with a 1 km floor. These tests need a trusted reference and are skipped where none is available. The primary Hispan-

iola configuration uses all expert coordinates, including records held out from threshold selection. The independent sensitivity run instead builds the reference from GeoMend's own first-pass output and uses no expert coordinates.

3. Locality-text consistency. The locality is parsed and the coordinate is flagged if it lies more than 3 km from where forward geodesy places the parsed offset, if the locality is a bare place name but the coordinate is more than 2 km from that place's town centre, if the named place is absent from the gazetteer, or if the coordinate is more than 5 km from the median of all coordinates sharing the same locality string.
4. Numerical feasibility. Flags latitude and longitude both rounded to two or fewer decimal places, or a location outside the user-provided bounding box.

### S2. Correction, road-walk, and georeferencing detail

Correction strategies are applied in order: consensus snapping among records that share an institutional locality identifier or the same locality string, repair of an unambiguous axis sign-flip, the road-distance walk for a stated offset, and nearest-road snapping otherwise. Locality parsing recognizes a numeric distance or distance range, miles or kilometres, an eight-point compass direction, and the named anchor before or after the offset phrase. Distance ranges are represented by their midpoint. The anchor is matched in GeoNames within the stated province where available.

For a parsed offset, Dijkstra traversal begins at the road node nearest the anchor and searches up to 1.8 times the stated distance. Candidate destinations must match the stated bearing within 55 degrees. Where several road branches qualify, a correction uses the candidate nearest the original coordinate, whereas a coordinate-less record uses the candidate nearest its straight-line distance-and-bearing seed. Candidates outside the corresponding proximity gate are rejected. The selected path is interpolated at the exact stated along-road distance rather than rounded to a graph node. Traversal confidence is one minus the absolute mismatch between the candidate and stated distance divided by the larger of the stated distance and 1 km. Candidates below 0.7 are rejected. Each accepted road placement is then moved 50 m perpendicular to the centreline, onto the side indicated by the record's original coordinate or straight-line seed. If that anchor does not distinguish a side, the side with the gentler elevation gradient is used. Off-trail taxa, from a configured list or identified automatically as taxa whose records predominantly fall far from any road, are exempted from nearest-road snapping.

For coordinate-less records, the Tier 2 gazetteer radius is set by GeoNames feature class from the calibration-split error of each class: 38.8 km for a populated place, 17.7 km for a mountain or terrain feature, and 36.7 km for a hydrographic feature. These values are the smallest observed-error thresholds containing at least 90% of their 254, 205, and 21 calibration records, respectively. Upward rounding to 0.1 km prevents the reported radius from falling below the selected observation. No calibration record reaches the spot class, so its 15 km value is explicitly a provisional fallback rather than a calibrated radius. The synchronization check and recomputation are in `repro/calibrate.py` and `outputs/calibration/calibration.md`. Place-name matching tolerates spelling variants, accent and encoding noise, and Spanish homophones. For Tier 3, the radius is the geographic extent of the named province or country.

### S3. Acceptance criteria

Every placement must be geographically plausible, and is flagged rather than assigned if it lands in water. A correction to an existing coordinate is withheld, and the record left flagged, when confidence is below 0.7, when the move is less than 1 km (a GPS-jitter-scale nudge that cannot fix a real error but can shift the point into a different climate-raster cell), or when the proposed coordinate lands more than 250 m in elevation from the collector-recorded value.

### S4. Keep-with-uncertainty

A flagged record, or a coordinate-less record that no tier could place, that still carries a usable original coordinate is kept with an elevated uncertainty radius calibrated to its flag reason (7.3 to 15.4 km, the smallest observed-error threshold containing at least 90% of calibration records for each reason). The exception is the gross-error tail flagged by the elevation, spatial-outlier, or nearest-neighbour detectors, which is not promoted and stays flagged for expert review.

### S5. Global claim and validation arms

The global claim arm contains preserved-specimen records belonging to species with no latitude and longitude on any record in the pinned snapshot. A locality was considered usable when its trimmed text contained at least three characters. Every qualifying record was retained. Because these records have no coordinate by definition, accuracy cannot be measured directly. We therefore drew a separate validation arm of coordinate-bearing records, removed their coordinates before georeferencing, and measured the great-circle distance between the GeoMend placement and the recorded coordinate.

The donor pool differs from the claim arm in its taxonomic and locality-text composition, so unweighted agreement would not represent the backlog. Donors were post-stratified to the claim distribution using clade, presence of a parseable distance-and-direction offset, and locality-length class. Sparse clades and cells were successively collapsed to kingdom-level and coarser cells until each retained cell contained at least 50 donors. Within each cell, donors received the claim-to-donor frequency ratio, normalized to mean one. All global agreement summaries use these weights. This comparison is agreement rather than independent truth because some recorded donor coordinates may themselves have been georeferenced from the same locality text.

### S6. Additional calibration, benchmarking, and interpretation details

Twelve thresholds are calibrated directly against the expert coordinates without using the validation set. Six are summary statistics of the manual coordinates, and six are selected from accuracy-coverage curves on the calibration split. Four determine whether to accept a correction: overall confidence, elevation tolerance, road-traversal confidence, and maximum correction distance. Two determine whether distance from a road or species range should trigger a flag. The resulting overall confidence threshold is 0.7, the road-distance threshold is 0.10 km, and the range-distance threshold is 1.0 km. GeoMend's confidence score describes how well the available evidence supports a candidate correction. It is neither a probability nor a true-positive rate.

Sixteen thresholds represent physical quantities, properties of external data, or field-practice conventions. The elevation test allows a 150 m mismatch because both the 30 m elevation model and recorded elevations are approximate. A coordinate rounded to two decimal places lies on a grid about 1.1 km across, so that grid sets the record's resolution limit and triggers the precision flag. The road-walk searches up to 1.8 times the stated distance because measured road sinuosity is about 1.7. It accepts bearings within 55 degrees because collectors commonly use eight-point compass sectors, each 45 degrees wide. Finally, road-derived placements are moved 50 m into adjacent habitat, close to the 54 to 56 m median road offset observed in the expert coordinates. Table S1 lists the remaining constants and their bases.

The remaining ten thresholds are user choices, but the results are insensitive to their exact values. We selected values by inspecting the Hispaniola records, then varied each threshold independently by 50% above and below its default. Across these tests, flagging recall changed by at most 0.2 percentage points, the fraction of records retaining a usable coordinate changed by at most 1.5 percentage points, and correction accuracy was essentially unchanged (Table S2). These thresholds mostly control how GeoMend identifies off-road taxa. We expose them because a region with sparser maps or a more ambiguous gazetteer may require different settings.

Applying GeoMend in a new region therefore requires local recalibration, not wholesale retuning. We recommend retaining the ten insensitive thresholds at their defaults and re-measuring the twelve calibrated thresholds from whatever local expert coordinates are available. The packaged calibration script performs this step from any truth table.

The Nature Plants benchmark tests whether GeoMend generalizes beyond Hispaniola. It contains 5,000 occurrence localities from 72 countries, each with an expert-assigned coordinate treated here as ground truth (Xie et al., 2025). We ran GeoMend with the same default configuration used for Hispaniola and made no regional adjustments. We compared its output with three baselines: GPT-4o evaluated on the same localities (Xie et al., 2025), the GEOLocate service (Rios & Bart, n.d.), and the centroid of each record's stated county. For each method, we measured both coverage and the distance over the Earth's surface from its placement to the published coordinate.

The Hispaniola benchmark provides a stricter record-level comparison with GEOLocate. We submitted each record's locality, province, and country to the GEOLocate web service and compared its best-ranked point with the same expert coordinate used to score GeoMend. We ran GEOLocate both with and without its road and highway correction to test whether that option explained any difference. The service's public documentation did not identify the road dataset underlying that option, so we cannot assess whether road-network coverage contributed to the result. Every response is cached with the analysis code, so reproducing the benchmark does not require querying the service again.

A wrong coordinate is not the only thing wrong with an occurrence database, and the two problems are worth separating. Records also cluster near roads and other accessible places because that is where collectors could reach, and this sampling bias can distort mapped diversity and species distribution models (Daru et al., 2018; He et al., 2025; Hughes et al., 2021; Kadmon et al., 2004; Wilcox et al., 2026; Zizka et al., 2021). Correcting a coordinate moves a record to where the specimen was actually collected, which is what a model needs from that record, but it does not change which places were visited. Positional accuracy is the target here, and sampling bias remains a separate property of the data that a user must address with the tools built for it.

Large language models can georeference locality text with accuracy comparable to established services (Fernando et al., 2026; Xie et al., 2025). However, general models typically provide neither calibrated uncertainty nor abstention when evidence is insufficient. Both are established georeferencing requirements (Chapman & Wieczorek, 2020; Wieczorek et al., 2004; Xie et al., 2025). We therefore compare GeoMend with a language model and the standard service to assess accuracy and generalization, while treating reproducibility, uncertainty, and abstention as separate design criteria. We evaluate detection and placement against expert truth on Hispaniola (Section 3.1), test global generalization against existing tools (Section 3.2), and measure downstream range recovery and the occurrence data returned for under-sampled species (Section 3.3).

GeoMend is designed to make every result reproducible from its inputs and configuration. It is an open-source Python package with a command-line interface under the MIT licence, and the same input table and configuration always produce the same output. All thresholds reside in one human-readable configuration file. Separate scripts regenerate the calibration, sensitivity analysis, benchmarks, downstream evaluation, and figures reported here. The source, configuration, and required data are archived as described in the Data Availability Statement.

### Table S1. Every threshold and how it is set

**Table S1.** Every GeoMend threshold, its value, and how that value is set. The 38 thresholds of Table 1 comprise 12 calibrated from the manual coordinates, 16 fixed with a stated external basis, and 10 with no external anchor and shown insensitive in Table S2. Toggles, curated lists, structural constants, inactive settings, and analysis-only fields are excluded. Regenerated by `repro/calibrate.py`. `--check-synced` fails if any empirically supported configured radius differs from the calibration-split value. The spot-feature radius is the sole provisional fallback because no calibration record reaches that class.

| Parameter | Value | How set | Basis |
|---|---|---|---|
| `coarse_georef.unc` | {P:38.8, T:17.7, H:36.7, S:15.0} | calibrated | ≥90% containment by feature class · S is an unsupported fallback |
| `flagging.correcti` | 250.0 | calibrated | point where corrections stop helping (~12% beyond 250 m) |
| `flagging.knn_outl` | 1.0 | calibrated | multiplier x each species' own p95 kNN distance |

| Parameter | Value | How set | Basis |
|---|---|---|---|
| flagging.max_dis[illegible] | 1.0 | calibrated | highest-F1 value in the calibration-split outside-range sweep |
| flagging.max_dis[illegible] | 0.10 | calibrated | highest-F1 value in the calibration-split distance-from-road sweep |
| flagging.max_sna[illegible] | 3.0 | calibrated | ~p75 of the paired-truth inlier offset distribution |
| flagging.min_acc[illegible] | 0.7 | calibrated | calibration-split accepted-error/coverage sweep |
| flagging.spatial_[illegible] | 0.5 | calibrated | multiplier x each species' own p95 point-to-centroid distance |

| Parameter | Value | How set | Basis |
|---|---|---|---|
| keep_with_uncerta | {skipped_off_road_ not_road_mode:15 low_accept… | calibrated | ≥90% containment of original-vs-truth error by flag reason |
| locality_consensu | 1.0 | calibrated | within-locality truth spread ~0 km · dispersion gate |
| road_distance.ma› | 6.0 | calibrated | capped-move sweep (median 1.3->0.7 km, %helped 58->66) |
| road_distance.mi | 0.7 | calibrated | direct comparison with road snapping · at values >=0.7 the road walk is more accurate in approximately 65% of cases |

| Parameter | Value | How set | Basis |
|---|---|---|---|
| `coarse_georef.def` | 25.0 | fixed | fallback = mid of the calibrated per-class radii (25 km) |
| `flagging.knn_outl` | 1.0 | fixed | GPS/rounding noise scale: sub-1 km scatter is not a real outlier |
| `flagging.max_dist` | 2.0 | fixed | settlement radius: a bare town name should fall within ~2 km of its centroid |
| `flagging.max_elev` | 150.0 | fixed | SRTM +-30 m at ridges + imprecise recorded elevations (deliberately forgiving) |

| Parameter | Value | How set | Basis |
| --- | --- | --- | --- |
| flagging.max_loca | 3.0 | fixed | separates compound-numeric errors from normal road-curvature / straight-line slack |
| flagging.max_prec | 2 | fixed | 2 dp ~ 1.1 km: coords rounded this coarse are approximate placeholders |
| flagging.min_acce | 1.0 | fixed | climate-raster cell ~1 km: sub-cell nudges can't fix range error but shift SDM cells |

| Parameter | Value | How set | Basis |
|---|---|---|---|
| flagging.ocean_bu | 0.5 | fixed | coastline-polygon resolution: forgives GADM coastline imprecision so coastal records aren't mis-flagged |
| flagging.road_hab | 50.0 | fixed | field-practice habitat offset · expert median 54 to 56 m from major roads · offset ablation did not reduce accuracy |
| keep_with_uncerta | 15.0 | fixed | fallback = max of the calibrated per-reason radii (15 km) |

| Parameter | Value | How set | Basis |
|---|---|---|---|
| locality_consensu | 0.5 | fixed | GPS noise scale: don't churn records already within 0.5 km of consensus |
| road_distance.bea | 55.0 | fixed | labels use 8-point compass (45 deg sectors) · ~half-sector + curvature slack -> 55 deg |
| road_distance.mir | 1.0 | fixed | method boundary: below 1 km a perpendicular snap already suffices |
| road_distance.ove | 1.8 | fixed | measured road sinuosity ~1.7x -> 1.8x Dijkstra search radius |

| Parameter | Value | How set | Basis |
| --- | --- | --- | --- |
| `text_georef.confi` | 0.8 | fixed | fixed pass-through tag above the 0.7 accept gate, not fit |
| `text_georef.local` | 5.0 | fixed | 5 km usability line: donors spread wider than this name more than one place |
| `coarse_georef.max` | 25.0 | free | decline if same-name candidates spread farther than this (-> sensitivity) |
| `coarse_georef.pro` | 1.5 | free | reject a proximity match beyond mult x province radius (-> sensitivity) |

| Parameter | Value | How set | Basis |
|---|---|---|---|
| `flagging.max_dis`[illegible] | 5.0 | free | same-locality-string divergence tolerance (no external basis -> sensitivity) |
| `flagging.offroad_`[illegible] | 1.0 | free | off-road if nearest road farther than this (no external basis -> sensitivity) |
| `flagging.offroad_`[illegible] | 5.0 | free | cap on road-distance correction for off-road taxa (-> sensitivity) |
| `flagging.offroad_`[illegible] | 0.3 | free | taxon off-road if >= this frac of records off-road (no external basis -> sensitivity) |

| Parameter | Value | How set | Basis |
|---|---|---|---|
| road_distance.ma› | 8.0 | free | max place-to-road anchoring distance (no external basis -> sensitivity) |
| text_georef.max_a | 25.0 | free | bare-name ambiguity guard, the Tier-1b analogue of the coarse one (-> sensitivity) |
| text_georef.roadw | 12.0 | free | reject a walk displacing the seed farther than this (-> sensitivity) |
| text_georef.roadw | 8.0 | free | walked point must land within this of the straight-line seed (-> sensitivity) |

### Table S2. Sensitivity of the free parameters

**Table S2.** Sensitivity of the principal metrics to the ten parameters that have no external anchor. Each is varied to 50% above and below its reported value, one at a time, with every other parameter held at its default. Entries are the full range of each metric across those values. Baseline: flagging recall 0.8965, corrected-median 1.049 km, fraction improved 0.673, usable coverage 0.8654. Regenerated on the calibration split by `repro/sensitivity.py`.

| Paramete | Default | Flagging recall (pp) | Corrected median (km) | Fraction improved (pp) | Usable coverage (pp) |
|---|---|---|---|---|---|
| `flagging.` | 5 | 0.21 | 0.010 | 0.12 | 0.02 |
| `flagging.` | 1 | 0.00 | 0.020 | 0.26 | 1.23 |
| `flagging.` | 0.3 | 0.00 | 0.009 | 0.44 | 1.49 |
| `flagging.` | 5 | 0.00 | 0.018 | 0.54 | 0.06 |
| `road_dist` | 8 | 0.00 | 0.007 | 0.34 | 0.00 |
| `coarse_ge` | 25 | 0.00 | 0.000 | 0.00 | 0.00 |
| `coarse_ge` | 1.5 | 0.00 | 0.000 | 0.00 | 0.00 |
| `text_geor` | 8 | 0.00 | 0.062 | 0.00 | 0.00 |
| `text_geor` | 12 | 0.00 | 0.048 | 0.00 | 0.00 |
| `text_geor` | 25 | 0.00 | 0.017 | 0.00 | 0.25 |
| **maximur** | | **0.21** | **0.062** | **0.54** | **1.49** |

### Table S3. Coverage-at-precision disposition

Disposition of all 16,913 Hispaniola occurrence records under GeoMend, ordered best-precision-first. Every record reaches exactly one tier. Where expert truth exists we report that tier's median error and its containment, the fraction of records whose truth falls within the assigned uncertainty radius. An endorsed coordinate (tiers 0 to 3, recommended at its labelled precision) is available for 74.8%, a coordinate of some kind for 99.8%, and only 30 records (0.2%) name no locatable place and receive none. Built by `repro/coverage_tiers.py`.

| Tier | n | % | median error | assigned radius | containment |
|---|---|---|---|---|---|
| 0 · trusted original | 2,645 | 15.6 | 0.70 km | — | 93% ≤5 km |
| 1a · precise correction | 1,208 | 7.1 | 0.82 km | — | 90% ≤5 km |
| 1b · text-georefere | 1,349 | 8.0 | 1.31 km | — | 81% ≤5 km |
| kept with uncertainty | 3,706 | 21.9 | 1.55 km | 13 km | 92% |

| Tier | n | % | median error | assigned radius | containment |
|---|---|---|---|---|---|
| 2 · gazetteer centroid | 1,737 | 10.3 | 6.41 km | class-specific | 93% |
| 3 · province centroid | 293 | 1.7 | 27.0 km | ~40 km | 84% |
| 3 · country centroid | 1,712 | 10.1 | 92.6 km | ~175 km | 95% |
| flagged for review | 4,233 | 25.0 | 1.18 km | — | — |
| no coordinate | 30 | 0.2 | — | — | — |

### Table S4. Taxonomic sources used in curating the Hispaniola reptile records

Revisions and systematic treatments consulted when checking or updating species identifications for the reptile records (Section 2.2). These accompany the archived dataset.

| Group | Source |
|---|---|
| *Anolis* | Schwartz, A. 1974. An analysis of variation in the Hispaniolan giant anole, *Anolis ricordi* Dumeril and Bibron. *Bulletin of the Museum of Comparative Zoology* 146, 89 to 146. |
| *Anolis* | Schwartz, A. & Henderson, R. W. 1982. *Anolis cybotes* (Reptilia: Iguanidae): the eastern Hispaniolan populations. *Milwaukee Public Museum Contributions in Biology and Geology* 49, 1 to 8. |
| *Anolis* | Köhler, G. & Hedges, S. B. 2016. A revision of the green anoles of Hispaniola with description of eight new species (Reptilia, Squamata, Dactyloidae). *Novitates Caribaea* 9, 1 to 135. |

| Group | Source |
| --- | --- |
| *Anolis* | Mahler, D. L., Lambert, S. M., Geneva, A. J., Ng, J., Hedges, S. B., Losos, J. B. & Glor, R. E. 2016. Discovery of a giant chameleon-like lizard (*Anolis*) on Hispaniola and its significance to understanding replicated adaptive radiations. *The American Naturalist* 188, 357 to 364. |
| *Audantia* | Köhler, G., Zimmer, C., McGrath, K. & Hedges, S. B. 2019. A revision of the genus *Audantia* of Hispaniola with description of four new species (Reptilia: Squamata: Dactyloidae). *Novitates Caribaea* 14, 1 to 104. |
| *Amphisbaena* | Thomas, R. & Hedges, S. B. 2006. Two new species of *Amphisbaena* (Reptilia: Squamata: Amphisbaenidae) from the Tiburon Peninsula of Haiti. *Caribbean Journal of Science* 42, 208 to 219. |

| Group | Source |
| --- | --- |
| Celestinae | Landestoy, M. A., Schools, M. & Hedges, S. B. 2022. A new genus and species of Caribbean forest lizard (Diploglossidae, Celestinae) from southern Hispaniola. *Zootaxa* 5219, 201 to 226. |
| Celestinae | Schools, M. & Hedges, S. B. 2024. A new forest lizard fauna from Caribbean islands (Squamata, Diploglossidae, Celestinae). *Zootaxa* 5554, 1 to 306. |
| *Ialtris* | Schwartz, A. & Thomas, R. 1965. The genus *Darlingtonia* (Serpentes) in Hispaniola, including a new subspecies from the Dominican Republic. Breviora 229, 1 to 10. |
| *Pholidoscelis* | Schwartz, A. & Klinikowski, R. F. 1966. The *Ameiva* (Lacertilia, Teiidae) of Hispaniola. II. Geographic variation in *Ameiva chrysolaema* Cope. *Bulletin of the Museum of Comparative Zoology* 133, 425 to 487. |

| Group | Source |
| --- | --- |
| *Pholidoscelis* | Schwartz, A. 1967. The *Ameiva* (Lacertilia, Teiidae) of Hispaniola. III. *Ameiva taeniura* Cope. *Bulletin of the Museum of Comparative Zoology* 135, 345 to 375. |
| *Pholidoscelis* | Hower, L. M. & Hedges, S. B. 2003. Molecular phylogeny and biogeography of West Indian teiid lizards of the genus *Ameiva*. *Caribbean Journal of Science* 39, 298 to 306. |
| *Pholidoscelis* | Goicoechea, N., Frost, D. R., De la Riva, I., Pellegrino, K. C. M., Sites, J. W., Rodrigues, M. T. & Padial, J. M. 2016. Molecular systematics of teioid lizards (Teioidea/Gymnophthalmoidea: Squamata) based on the analysis of 48 loci under tree-alignment and similarity-alignment. *Cladistics* 32, 624 to 671. |

### Table S5. Detection under five species-reference sets

Three of GeoMend's detectors (`FAR_FROM_RANGE`, `SPATIAL_OUTLIER`, `KNN_OUTLIER`) screen a record against a per-species reference built from coordinates of its conspecifics. In the primary run that reference is the expert table, which is also the truth we score against, so the arms below progressively remove that dependence. Detection is measured on the 6,322 held-out records. Endorsed coverage is over all 16,913. Built by `repro/reference_independence.py`.

| Reference set | recall >5 km | recall >1 km | recall, reviewer flag | precision >1 km | endorsed coverage |
|---|---:|---:|---:|---:|---:|
| Full expert table (primary run) | 92.0% | 88.5% | 79.0% | 56.6% | 74.8% |
| Expert table, each record left out of its own reference | 92.2% | 89.0% | 80.0% | 56.3% | 73.9% |

| Reference set | recall >5 km | recall >1 km | recall, reviewer flag | precision >1 km | endorsed coverage |
|---|---|---|---|---|---|
| GeoMend own first-pass output, no expert data | 91.7% | 89.0% | 79.8% | 55.9% | 71.1% |
| Expert coordinates from training records only | 90.4% | 87.4% | 80.1% | 56.5% | 76.9% |
| No species-level reference at all | 88.0% | 85.0% | 73.6% | 57.3% | 84.4% |

| Reference set | recall >5 km | recall >1 km | recall, reviewer flag | precision >1 km | endorsed coverage |
|---|---|---|---|---|---|
| Coordinat for reference | 27% | 19% | 10% | 59% | not applicable |

Weakening the reference costs recall and, because fewer records are flagged, raises endorsed coverage. The two move in opposite directions, so the arms bracket the dependence rather than replacing one number with another. The worst case, no species-level reference at all, still recovers 88% of coordinates more than 5 km from truth.

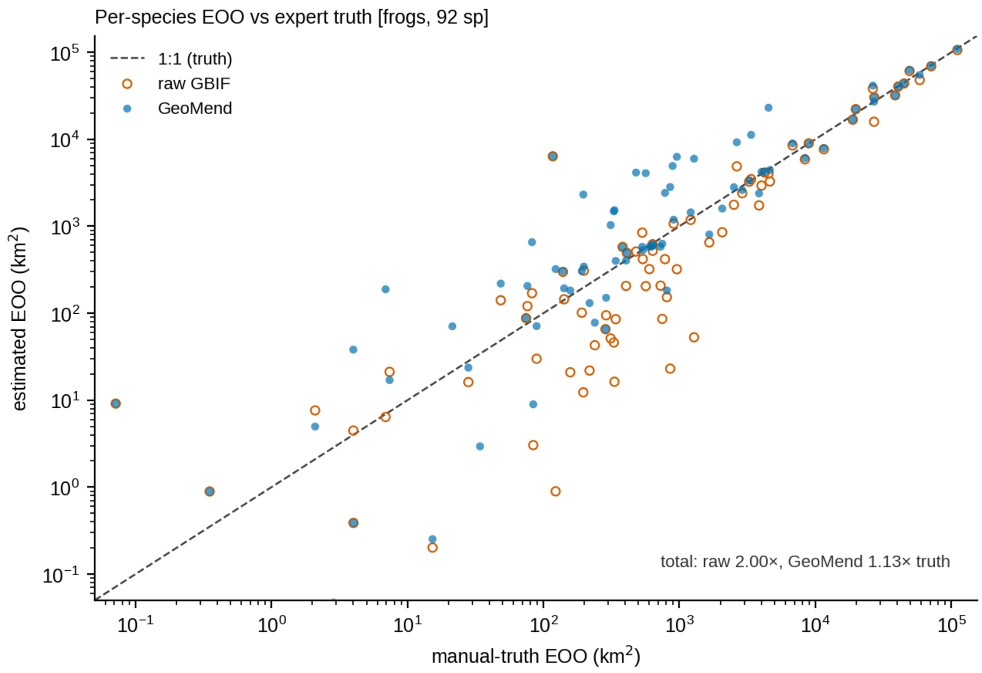


**Figure S1. Per-species extent of occurrence recovery for frogs.** Raw and GeoMend extents of occurrence are compared with expert truth for the 94 species with expert EOO, of which 81 are measurable in all three coordinate sets.

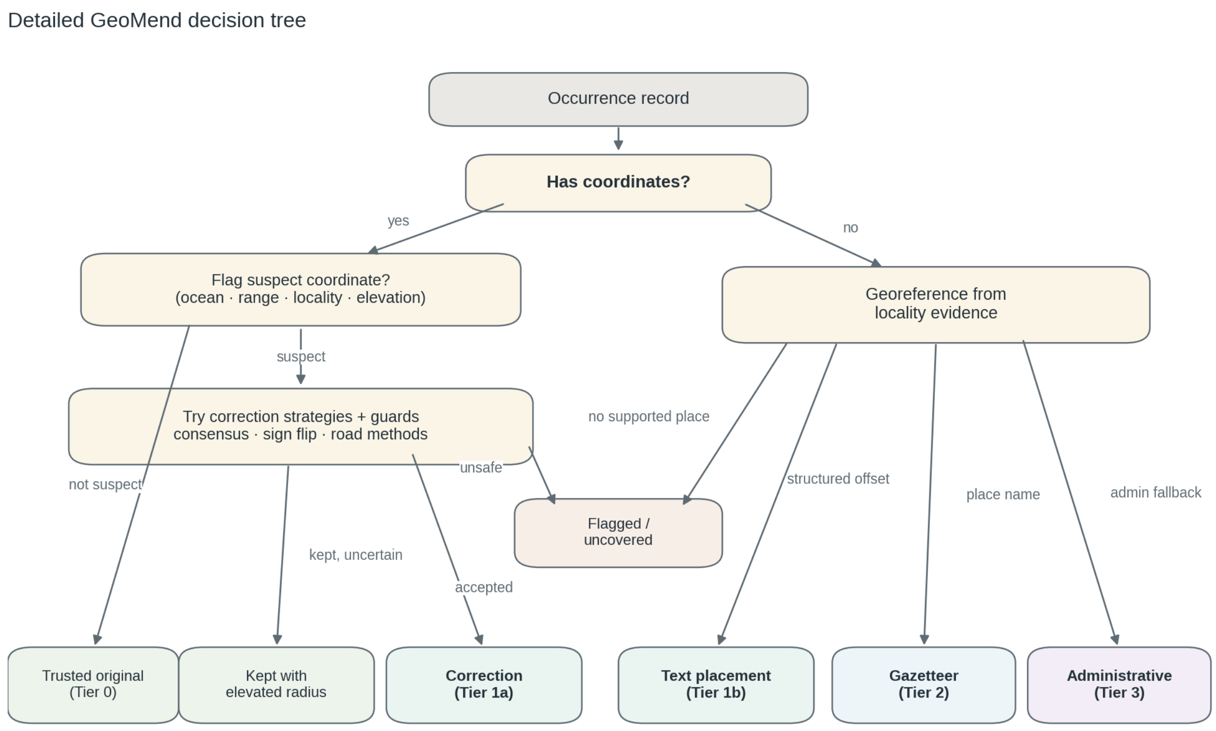


**Figure S2. Detailed GeoMend decision tree.** Coordinate-bearing records are screened before ordered correction strategies are attempted, and records without coordinates are routed through structured-text, gazetteer, and administrative placements. Failed or unsupported placements remain flagged or uncovered rather than being reported as endorsed coordinates.

Chapman, A. D., & Wieczorek, J. R. (2020). *Georeferencing best practices*. GBIF Secretariat. https://doi.org/10.15468/doc-gg7h-s853

Daru, B. H., Park, D. S., Primack, R. B., Willis, C. G., Barrington, D. S., Whitfeld, T. J. S., Seidler, T. G., Sweeney, P. W., Foster, D. R., Ellison, A. M., & Davis, C. C. (2018). Widespread sampling biases in herbaria revealed from large-scale digitization. *New Phytologist*, *217*(2), 939–955. https://doi.org/10.1111/nph.14855

Fernando, A., Ranathunga, S., Stock, K., Prasanna, R., & Jones, C. B. (2026). Georeferencing complex relative locality descriptions with large language models. *International Journal of Geographical Information Science*, 1–33. https://doi.org/10.1080/13658816.2026.2613291

He, J., Yang, W., You, Q., Hu, Q., Cong, M., Tian, C., & Ma, K. (2025). Impacts of road networks on the geography of floristic collections in China. *Plant Diversity*, *47*(3), 403–414. https://doi.org/10.1016/j.pld.2025.02.001

Hughes, A. C., Orr, M. C., Ma, K., Costello, M. J., Waller, J., Provoost, P., Yang, Q., Zhu, C., & Qiao, H. (2021). Sampling biases shape our view of the natural world. *Ecography*, *44*, 1259–1269. https://doi.org/10.1111/ecog.05926

Kadmon, R., Farber, O., & Danin, A. (2004). Effect of roadside bias on the accuracy of predictive maps produced by bioclimatic models. *Ecological Applications*, *14*(2), 401–413. https://doi.org/10.1890/02-5364

Rios, N. E., & Bart, H. L. (n.d.). *GEOLocate: A platform for georeferencing natural history collections data*. Tulane University Biodiversity Research Institute; https://www.geo-locate.org.

Wieczorek, J., Guo, Q., & Hijmans, R. J. (2004). The point-radius method for georeferencing locality descriptions and calculating associated uncertainty. *International Journal of Geographical Information Science*, *18*(8), 745–767. https://doi.org/10.1080/13658810412331280211

Wilcox, R. C., Baniaga, A. E., Hill, A. P., Young, A., Johnson, R. F., & Jacobs, S. J. (2026). Documenting biodiversity with digital data: Comparing and contrasting the efficacy of specimen-based and observation-based approaches. *New Phytologist*, *251*, 721–736. https://doi.org/10.1111/nph.70406

Xie, Y., Park, D. S., Sinnott-Armstrong, M. A., Ho, J., Chen, T., Weakley, A. S., Aguirre Lopez, L. J., Choi, J., Laitinen, M. M., Steeves, N. A., Huang,

C. H., Xu, R., & Feng, X. (2025). Using large language models to address the bottleneck of georeferencing natural history collections. *Nature Plants*, *11*(12), 2446–2450. https://doi.org/10.1038/s41477-025-02162-y

Zizka, A., Antonelli, A., & Silvestro, D. (2021). sampbias, a method for quantifying geographic sampling biases in species distribution data. *Ecography*, *44*, 25–32. https://doi.org/10.1111/ecog.05102
